# Transmembrane polar relay drives the allosteric regulation for ABCG5/G8 sterol transporter

**DOI:** 10.1101/2020.10.06.327825

**Authors:** Bala M. Xavier, Aiman A. Zein, Angelica Venes, Junmei Wang, Jyh-Yeuan Lee

**Author notes:** Contributed equally (Bala M. Xavier, Aiman A. Zein). Department of Cellular and Molecular Medicine, Faculty of Medicine, University of Ottawa, ON, Canada. Correspondence and; (J.W.) (J.-Y.L.).

## Abstract

The heterodimeric ATP-binding cassette (ABC) sterol transporter, ABCG5/G8, is responsible for the biliary and transintestinal secretion of cholesterol and dietary plant sterols. Missense mutations of ABCG5/G8 can cause sitosterolemia, a loss-of-function disorder characterized by plant sterol accumulation and premature atherosclerosis. A new molecular framework was recently established by a crystal structure of human ABCG5/G8 and reveals a network of polar and charged amino acids in the core of the transmembrane domains, namely polar relay. In this study, we utilize genetic variants to dissect the mechanistic role of this transmembrane polar relay in controlling ABCG5/G8 function. We demonstrated a sterol-coupled ATPase activity of ABCG5/G8 by cholesteryl hemisuccinate (CHS), a relatively water-soluble cholesterol memetic, and characterized CHS-coupled ATPase activity of three loss-of-function missense variants, R543S, E146Q, and A540F, which are respectively within, in contact with, and distant from the polar relay. The results established an *in vitro* phenotype of the loss-of-function and missense mutations of ABCG5/G8, showing significantly impaired ATPase activity and loss of energy sufficient to weaken the signal transmission from the transmembrane domains. Our data provide a biochemical evidence underlying the importance of the polar relay and its network in regulating the catalytic activity of ABCG5/G8 sterol transporter.

## 1. Introduction

All living cells depend on the ability to translocate nutrients, metabolites, and other molecules across their membranes. One major way to achieve this is through membrane-anchored transporter proteins. The evolutionarily conserved ATP-binding cassette (ABC) transporter superfamily, for example, carry out ATP-dependent and active transport of a wide range of substances across cellular membranes, including both hydrophilic and hydrophobic molecules such as sugars, peptides, antibiotics, or cholesterol [1–4]. As a key component of cellular membranes, cholesterol constitutes ~50% of cellular lipid content; it is also the precursor of steroid hormones that modulate gene regulation and bile acids that enable nutrient absorption. Translocation of cholesterol molecules on biological membranes plays an essential role in maintaining cellular and whole-body cholesterol homeostasis. Thus, excess cholesterol needs to be eliminated from cells and tissues through either sterol acceptors in the circulation or direct excretion into the bile or the gut [5,6]. A large body of evidence indicates that ABC sterol transporters regulate cholesterol metabolism, and their defects are associated with dysregulation of whole-body cholesterol homeostasis, a major risk factor for cardiovascular diseases [7,8]. Yet we have almost no understanding of how these transporters actually translocate cholesterol molecules and how the sterol-transport process is controlled by ATP catalysis. Given the dysregulation of cholesterol metabolism as a major risk factor for cardiovascular disease, there is a pressing need to elucidate of mechanism of these transporters in moving molecules across the cell membranes.

Recent progress in solving a heterodimeric crystal structure of human ABCG5 and ABCG8 establishes a new molecular framework towards such a mechanistic understanding of ABC sterol transporters. ABCG5 and ABCG8 are half-sized ABC sterol transporters and co-expressed on the apical surface of the hepatocytes along the bile ducts and the enterocytes from the intestinal brush-boarder membranes [9,10]. ABCG5 and ABCG8 function as obligate heterodimers (ABCG5/G8) and serve as the primary and indispensable sterol-efflux pump that effectively exports excess cholesterol, non-cholesterol sterols, and dietary plant sterols into the bile and the intestinal lumen. In mammals, most cholesterol is eliminated by its metabolism into bile acids or via biliary secretion as free cholesterol. The latter is considered as the last step of reverse cholesterol transport (RCT), where ABCG5/G8 accounts for more than 75% biliary cholesterol secretion [11–14]. Recent studies have shown that in human subjects and animal models, ABCG5/G8 is also responsible for eliminating neutral sterols by the transintestinal cholesterol efflux (TICE), a cholesterol-lowering process independent of RCT [15]. Physiologically, ABCG5/G8 thus plays an essential role in controlling cholesterol homeostasis in our bodies.

In general, the smallest functional unit of an ABC transporter consists of two transmembrane domains (TMD1 and TMD2) and two nucleotide-binding domains (NBD1 and NBD2), and both NBDs concertedly bind and hydrolyze ATP to provide the energy and drive substrate transport. The TMDs, on the other hand, have shown to share low sequence similarity in the amino acid sequences and three-dimensional structural folds, suggesting substrate-specific mechanisms for individual transporters [16]. Mechanistic analyses of ABC cholesterol transporters have largely centered on sequence requirement at the canonical ATP-binding sites [17–20], whereas limited is known about sterol-protein interaction and its relationship with the ATP catalysis. Recent progress solving a crystal structure of human ABCG5/G8 revealed a unique TMD fold and several structural motifs [21]. Of particular, for each subunit, a network of polar and charged amino acids is present in the core of the TMD, namely polar relay, whose role remains to be characterized. A triple-helical bundle is located at the transmission interface between the NBD and the TMD and consists of an elbow connecting helix, a hot-spot helix (also known as E-helix), and an intracellular loop-1 (ICL1) coupling helix. However, on the triple-helical bundle or the transmembrane polar relay, several residues have been shown to bear disease-causing missense mutations from sitosterolemia or other metabolic disorders with lipid phenotypes (**Figure 1A**). Notably, several disease-causing mutations are clustered in the membrane-spanning region or at the NBD-TMD interface [8,22]. This suggests the unique roles of these structural motifs in regulating the ABCG5/G8 function; yet, no prior knowledge was available to explain the role of these structural motifs in the sterol-transport function.

**Figure 1.**
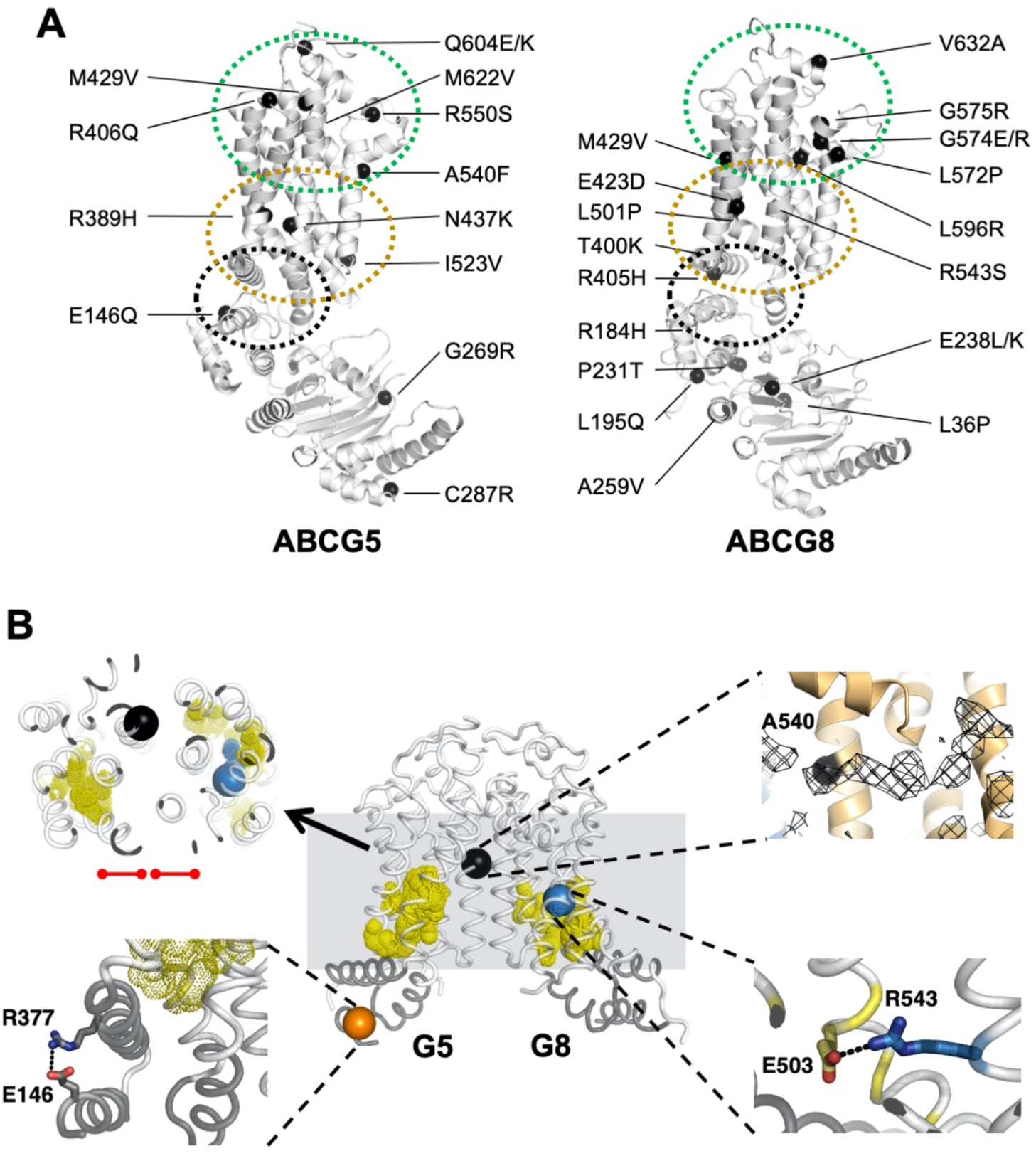
Disease-causing mutations and SNPs in ABCG5/G8. **A**, Localization of ABCG5/G8 residues carrying missense mutations. The positions of disorder-related polymorphisms or mutations are highlighted in black spheres on the structures of ABCG5 (PDB ID: 5D07, chain C) and ABCG8 (PDB ID 5D07, chain D). Structural motifs are indicated in dashed ovals: triple-helical bundle (black), TMD polar relay (yellow), and extracellular domain with re-entry helices (green). **B**, Microenvironment of G5-E146, G5-A540, and G8-R543. *(Middle)* The transmembrane domains (white) and the triple helical bundle (grey) are plotted in tube-styled cartoon presentation, showing the a-carbons (spheres) of G5-E146 (orange), G8-R543 (blue), and G5-A540 (black). The polar relays are plotted in dotted yellow spheres. *(Top-left)* Slapped top view shows G5-A540 situated more than 10Å away from the polar relay of either subunit (red dot-ended lines). *(Top-right)* Near G5-A540 shows a cholesterol-shaped electron density (mesh) in the crystal structure of ABCG5/G8. Fo-Fc difference electron density map was contoured at 3.0s. *(Bottom-left)* At the triple helical bundle of ABCG5, E146 interacts with R377 through their side-chain termini in a distance of hydrogen bonding, 3.5Å (black dashed line). *(Bottom-right)* In ABCG8 polar relay, R543 interacts E503 through their side-chain termini in a distance of hydrogen bonding, 3.1Å (black dashed line).

Loss-of-function (LOF) mutations in ABCG5 or ABCG8 are linked to sitosterolemia, a rare autosomal recessive disease, while several other missense mutations are also associated with other lipid disorders, such as gallstone formation or elevated LDL cholesterol [23–28]. At the cellular level, many of the missense mutations lead to defects in post-translational trafficking of ABCG5/G8 from the endoplasmic reticulum (ER), an abnormality commonly observed in other ABC transporters with missense mutations, *e.g.,*ΔF508 mutation in the cystic fibrosis transmembrane conductance regulator (CFTR or ABCC7) [29,30]. However, specific missense mutants of ABCG5/G8 heterodimers have shown no defect in protein maturation [29], suggesting alternative disease-causing mechanisms. Therefore, studies of these mutants will not only show how they alter the transporter activity, but will also provide mechanistic insights into the function of wild-type (WT) ABCG5/G8 sterol transporter.

Disease mutations are instrumental in studying the mechanisms of affected proteins *in vitro, e.g.*, familial hypercholesterolemia mutations for proteins involved in low-density lipoprotein metabolism [31]. Guided by the structural framework of ABCG5/G8, we can now investigate its mechanisms using enzymological approaches with purified proteins. For this, we first need to establish at least one robust and consistent *in vitro* functional assay. Using ATPase activity as the functional benchmark in this study, we have optimized an *in vitro* colorimetric ATPase assay that allows high-throughput activity assessment of detergent-purified ABCG5/G8. Using a soluble cholesterol memetic, cholesteryl hemisuccinate (CHS), we report here the CHS-stimulated ATP hydrolysis by ABCG5/G8 proteo-micelles, consisting of phospholipids, cholate, and dodecylmaltoside (DDM), and present an enzymatic analysis for the sterol-coupled ATPase activity on ABCG5/G8 sterol transporter. Using ATPase activity as functional readout of ABCG5/G8, we show differentially inhibition of the CHS-stimulated ATPase activity by three LOF missense mutants, two sitosterolemia mutations and one sterol-binding mutation, where residues bearing the two disease mutations are located along the polar relay. Our data hereby demonstrate the mechanistic basis on regulating ABCG5/G8 function by the transmembrane polar relay (**Figure 1B**).

## 2. Results

### 2.1. CHS stimulates ATP hydrolysis by wild-type (WT) ABCG5/G8

Despite the known physiological role of ABCG5/G8 in biliary and intestinal cholesterol secretion, only indirect evidence of sterol-coupled transporter activity was detected by using steroid mimetics, such as androstan or bile acids [32,33]. In this study, we investigated a direct sterol-coupled ATPase activity by using CHS, a cholesterol mimetic that is more soluble in aqueous solution. First, to overcome low sensitivity of detecting the ABCG5/G8 ATPase activity by previous protocols, we have optimized the ATPase assay for ABCG5/G8 by adopting a previous assay [34] and a colorimetric bismuth citrate-based detection approach [35]. As described and explained in Materials and Methods, this optimized assay significantly reduces the background noise due to cloudiness by phospholipid/cholate/DDM mixtures, which improves the detecting sensitivity of liberated inorganic phosphate within the first few minutes and allows us to calculate more accurate rates of ATP hydrolysis. We show here that CHS can significantly stimulate ABCG5/G8-mediated ATP hydrolysis when co-incubated with sodium cholate (a bile acid) and *E. coli* polar lipids (**Figure 2**). Using 5 mM of ATP, the basal activity of ABCG5/G8 was calculated as 160 ± 15 nmol/min/mg (n = 4), similar to reported values, whereas in the presence of CHS, the specific ATPase activity of ABCG5/G8 reached 565 ± 30 nmol/min/mg (n = 8), three-four times higher than that in the absence of CHS (**Figures 3A & 3B**). Absence of cholate was unable to activate the ATP hydrolysis, consistent with the previous studies (data not shown) [33]. In addition, the activity was inhibited either by orthovanadate, an ATPase inhibitor [36] (**Figure 3C**), or by a catalytically-deficient mutant ABCG5_WT_/G8_G2I6D_ (G8-G216D) [18], which displayed no ATP hydrolysis (**Figures 3A & 3B**). The specific activity of ATP hydrolysis by ABCG5/G8 is by far the highest in comparison with the previously reported values [33].

**Figure 2.**
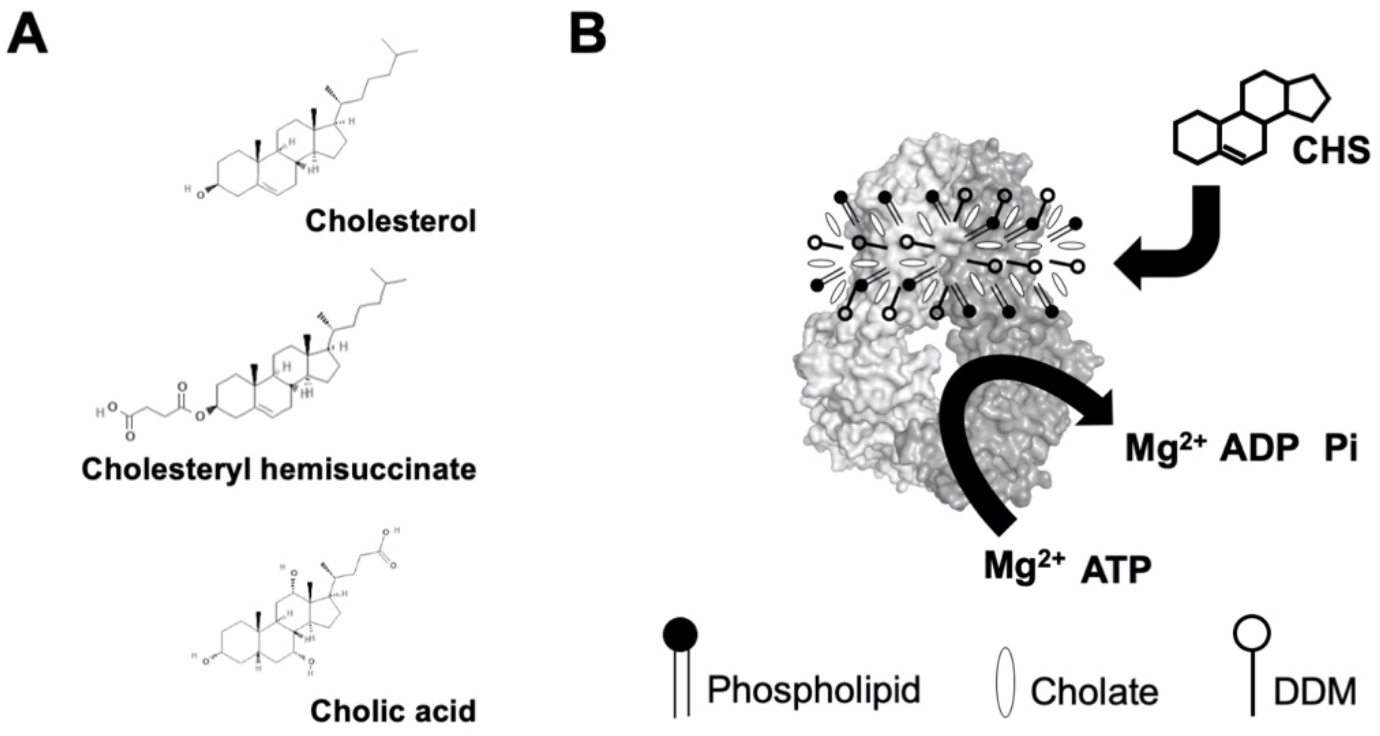
**A**, Chemical structures of cholesterol, cholesteryl hemisuccinate (CHS) and cholic acid (cholate). Source: PubChem. **B**, Schematic illustration of sterol-coupled ATPase activity of ABCG5/G8. DDM-purified ABCG5/G8 (light/dark grey surface) is preincubated with phospholipids and cholate. Addition of CHS (four-ringed steroid structure) stimulates hydrolysis of ATP to ADP and inorganic phosphate (Pi) in the presence of the divalent magnesium ions (Mg^2+^). Using the colorimetric and bismuth citrate-based assay, the liberated Pi is then captured by ammonium molybdate in the presence ascorbic acid. The color is developed upon mixing with bismuth citrate and sodium citrate, and the absorbance was measured at 695 nm. See details in Materials and Methods.

**Figure 3.**
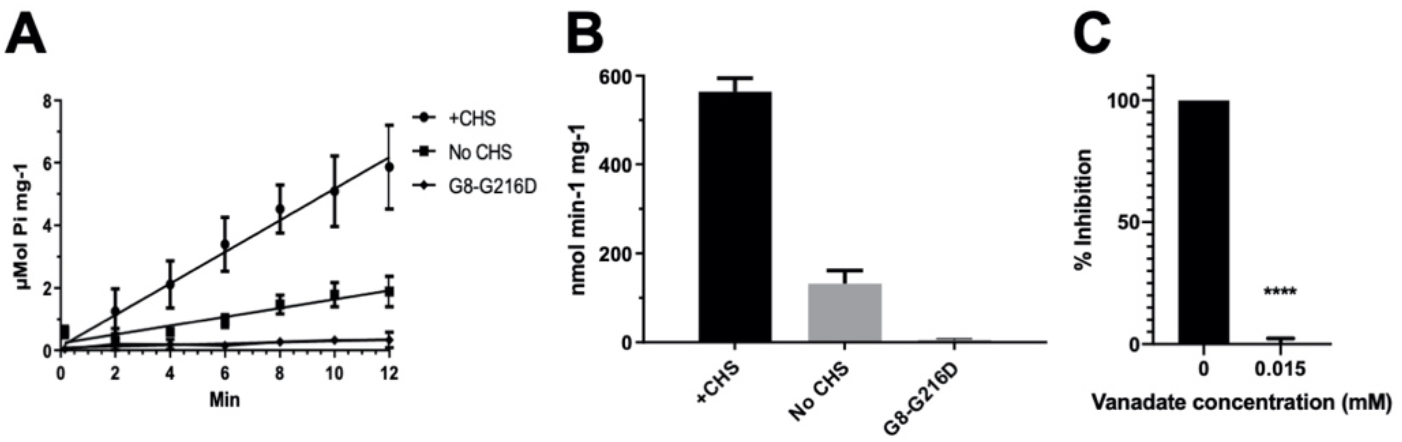
ATPase activity of ABCG5/G8. The ATP hydrolysis was used as a measure of ABCG5/G8 ATPase activity at 37°C in a condition with 5mM ATP and 4.1mM CHS. The protocol is entailed in Materials and Methods. **A**, Data points are presented as the means ± standard deviations from four-to-eight independent experiments using two-to-four independently purified proteins, where not visible, the error bars are covered by the plot symbols. A linear regression, plotted from the first 12 minutes, is used to calculate the specific activities. **B**, Bar graphs show the specific activities of ATP hydrolysis by WT in the presence and absence of CHS and the catalytically deficient mutant G8-G216D in the presence of CHS. The specific activity of WT in the absence of CHS is regarded as the basal ABCG5/G8 ATPase activity. **C**, Bar graphs represent percentage inhibition of ABCG5/G8 ATPase activity by 0.015mM orthovanadate, a P-value of <0.0001 obtained using ordinary one-way ANOVA (Prism 8).

### 2.2. The lipid environments fine-tune ABCG5/G8 ATPase activity

ABC transporters need to function in phospholipid-embedded environment. However, it is unknown whether the ABCG5/G8 function is controlled by phospholipids of specific headgroups or in specific lipid compositions. Because a high concentration of bile acids is required to activate ABCG5/G8 ATPase activity, attempts to use reconstituted proteoliposomes failed due to the immediate solubilization of the reconstituted proteins. To facilitate the assessment of mutant functions, we evaluated the lipid environments to obtain the most optimal assay conditions. To study the effect of lipid conditions and phospholipid species on the ABCG5/G8 function, we analyzed the CHS-coupled ATPase activity in the presence of two polar lipid extracts under a condition of fixed concentrations of sodium cholate and CHS (Materials and Methods). Using *E. coli* polar lipids, we carried out ATP concentration-dependent ATPase assay to determine the Michaelis-Menten kinetic parameters of CHS-stimulated ATP hydrolysis. We have observed the maximal ATP hydrolysis by ABCG5/G8 at concentrations slightly over 2.5 mM of ATP with a V_max_ of 677.1 ± 25.6 nmol/min/mg, a K_M_(ATP) of 0.60 mM, and a k_cat_ of 1.69 s^−1^. When using bovine liver polar lipids, we observed ~3.5-fold lower catalytic rate of ATP hydrolysis and ~50% higher K_M_(ATP) (**Figure 4A & Table 1**). In the current study, polar lipids, cholate (bile acid) and CHS were all present in the reaction, indicating that the presence of *E. coli* polar lipids results in higher ATP association and consequently better stimulates ABCG5/G8 ATPase activity. When comparing the calculated values of k_cat_ and k_cat_/K_M_, we indeed observed an overall 5-fold higher turnover rate in the presence of *E. coli* polar lipids than liver polar lipids (**Table 1**).

**Figure 4.**
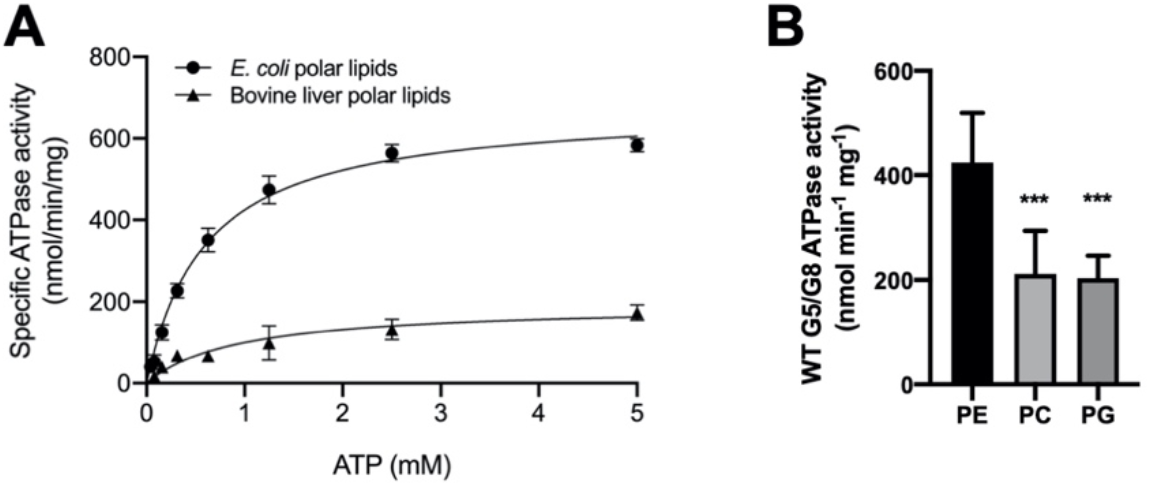
Lipid dependence of ABCG5/G8 ATPase activity. **A**, Purified ABCG5/G8 was assayed in the presence of either *E. coli* or bovine liver polar lipids, and the specific activities of ATP hydrolysis were obtained by the ATP concentration-dependent experiments (0-5mM ATP). Both curves are fitted to the Michaelis-Menten equation (Prism 8), and using two independently purified proteins, the means of at least three independent experiments along with standard deviations are plotted here. The kinetic parameters are listed in Table 1. **B**, In a condition of 5mM ATP and 4.1mM CHS, ATP hydrolysis of purified ABCG5/G8 was assayed in the presence of egg PE, soy PC or egg PG, a P-value of 0.0006 and 0.0003, respectively, obtained using ordinary one-way ANOVA (Prism 8).

**Table 1.**
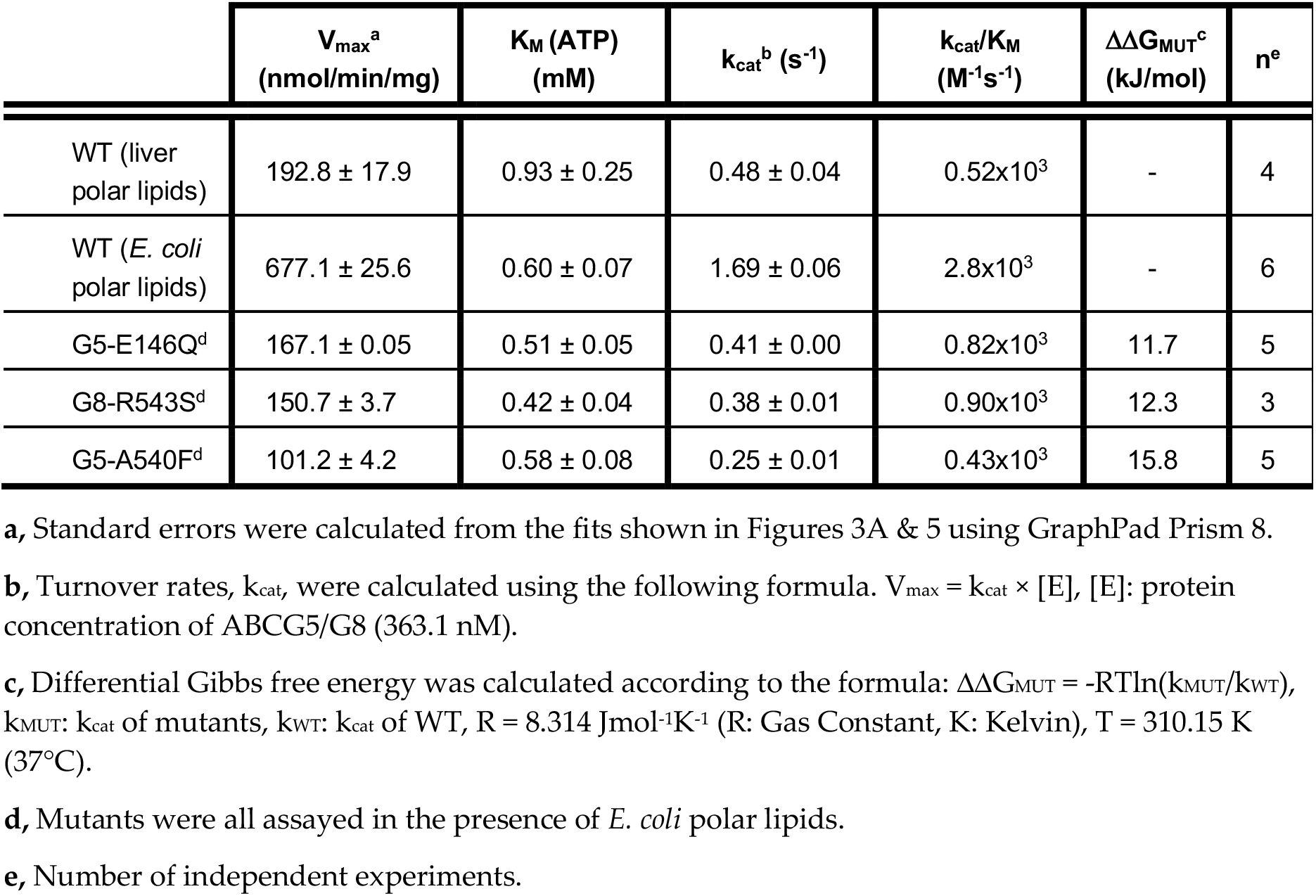
Dependence of ABCG5/G8 ATPase Activity on ATP.

To determine the dependence of phospholipid headgroups, we tested three most abundant phospholipids in either lipid extract on the ATP hydrolysis by ABCG5/G8, *i.e.*, phosphatidylethanolamine (PE), phosphatidylcholine (PC), and phosphatidylglycerol (PG) (see Materials and Methods). Preincubation with egg PE resulted in the highest specific activity, while the use of soy PC or egg PG only led to slightly higher ATP hydrolysis than the basal activity (**Figure 4B**). Interestingly, PE, the phospholipid found in both *E. coli* and liver lipids, is sufficient to stimulate ATP hydrolysis in ABCG5/G8 to almost the highest specific activity, as reported here. In the meantime, using PC or PG alone, the specific activity of ABCG5/G8 was also higher than that obtained with the liver polar lipid mixture. These results suggest phospholipid headgroups in regulating the ABCG5/G8 ATPase activity. Further investigations are necessary to pinpoint the effects of individual types of phospholipids on the sterol transporter function.

### 2.3. Missense mutants impair CHS-coupled ATPase Activity of ABCG5/G8

Using the CHS-coupled ATPase activity as the functional readout, we initiated studies in the catalytic mechanism of ABCG5/G8 by exploiting the transporter’s missense mutations that undergo proper trafficking to post-ER cell membranes (ER-escaped mutants). In this study, we have used *Pichia pastoris* yeast and expressed recombinant proteins of G8-G216D, a catalytically deficient mutant [18], ABCG5_EI46Q_/G8_WT_(G5-E146Q) and ABCG5_WT_/G8_R543S_ (G8-R543S), two loss-of-function/sitosterolemia missense mutants [22,37], and ABCG5_A54OF_/G8_WT_ (G5-A540F), a loss-of-function mutant with putative sterol-binding defect [21] (**Figure 1B & Supplementary Figure 1**).The purified mutants were preincubated with *E. coli* polar lipids and sodium cholate as described above. As shown in **Figure 5**, when compared with WT, the sitosterolemia missense mutants, G5-E146Q and G8-R543S, show a ~80% reduction of the specific activity in CHS-coupled ATP hydrolysis (160±15 nmol/min/mg and 150 ± 5 nmol/min/mg, respectively). The sterol-binding mutant G5-A540F, when compared to WT, shows a ~90% reduction of the specific activity in CHS-coupled ATP hydrolysis (90 ± 10 nmol/min/mg). Similar levels of activity reduction were also observed for non-CHS-coupled ATP hydrolysis (**Supplementary Figure 2**). We then performed ATP concentration-dependent experiments and analyzed the Michaelis-Menten kinetics for these three mutants. For all mutants, K_M_(ATP) remained nearly the same as compared to WT, but the mutants displayed a 40-60% reduction in the catalytic rate (**Table 1**). This result suggests that the mutants do not alter their ability of the nucleotide association, and other molecular events contribute to the reduction of the specific ATPase activity.

**Figure 5.**
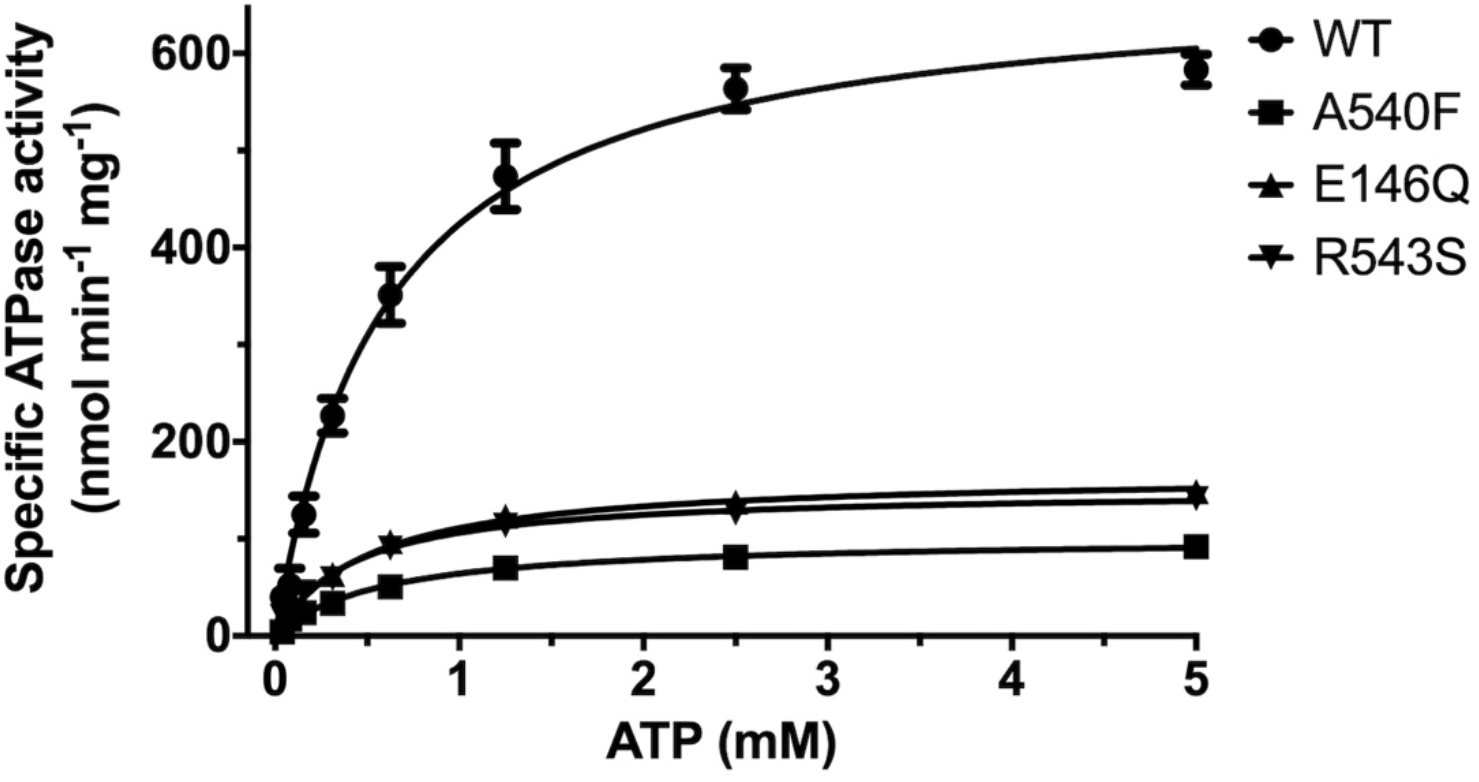
ATP-Dependence of ABCG5/G8 ATPase Activity. Purified proteins were assayed in the presence of *E. coli* polar lipids, and the specific activities of ATP hydrolysis were obtained by the ATP concentration-dependent experiments (0-5mM ATP). The curves are fitted to the Michaelis-Menten equation (Prism 8), and using two-to-four independently purified proteins, the means of at least three independent experiments along with standard deviations are plotted here. The kinetic parameters are listed in Table 1.

The effects of CHS on ABCG5/G8 WT and mutants were further investigated by measuring the ATP hydrolysis in the CHS concentration-dependent manner at a saturated ATP concentration (5mM here). Purified proteins were preincubated with *E. coli* polar lipids, sodium cholate and a wide range of CHS concentrations (0.064mM - 4.1mM). For WT, we obtained a V_max_ of 702.9 ± 50.7 nmol/min/mg, a K_M_(CHS) of 0.79 mM, and a k_cat_ of 1.74 s^−1^ (**Figure 6 & Table 2**). In the presence of *E. coli* polar lipids, the catalytic rates were similar between the CHS and ATP-dependent ATPase activities, a V_max_ of ~700 nmol/min/mg, which is about four times higher than that in the presence of liver polar lipids (**Tables 1 & 2**) and more than two-fold higher than the previously reported value, ~290 nmol/min/mg [33]. The catalytic rates of the mutants decreased by 70-90%, but except for G5-A540F, both G5-E146Q and G8-R543S displayed significantly larger K_M_(CHS), up to two-fold increase. This suggests a more profound impact of sitosterolemia mutations on the ABCG5/G8 ATPase activity through sterol-protein interaction or structural changes.

**Figure 6.**
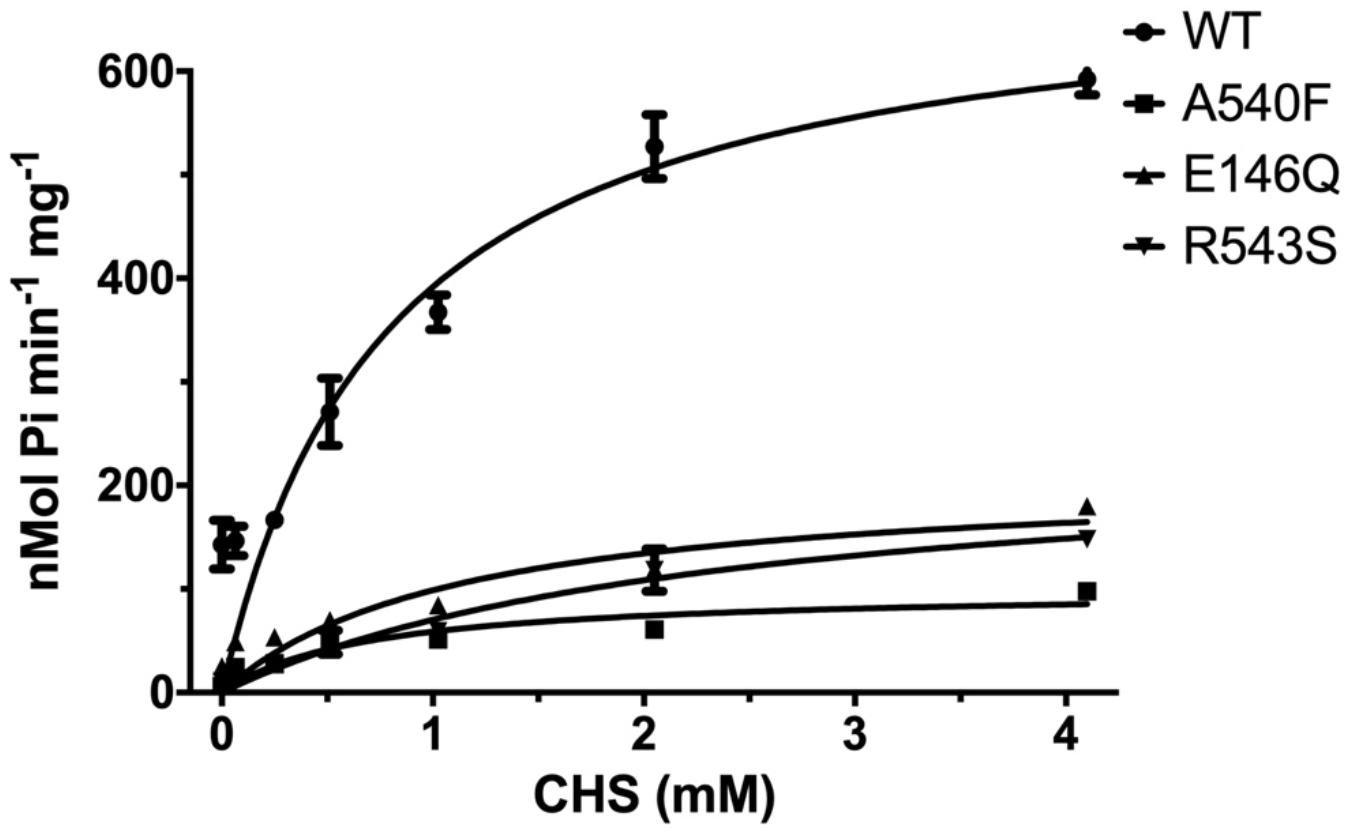
CHS-Dependence of ABCG5/G8 ATPase Activity. Purified proteins were assayed in the presence of *E. coli* polar lipids, and the specific activities of ATP hydrolysis were obtained by the CHS concentration-dependent experiments (0-4.1mM CHS). The curves are fitted to the Michaelis-Menten equation (Prism 8), and using two independently purified proteins, the means of at least two independent experiments along with standard deviations are plotted here. The kinetic parameters are listed in Table 2.

**Table 2.**
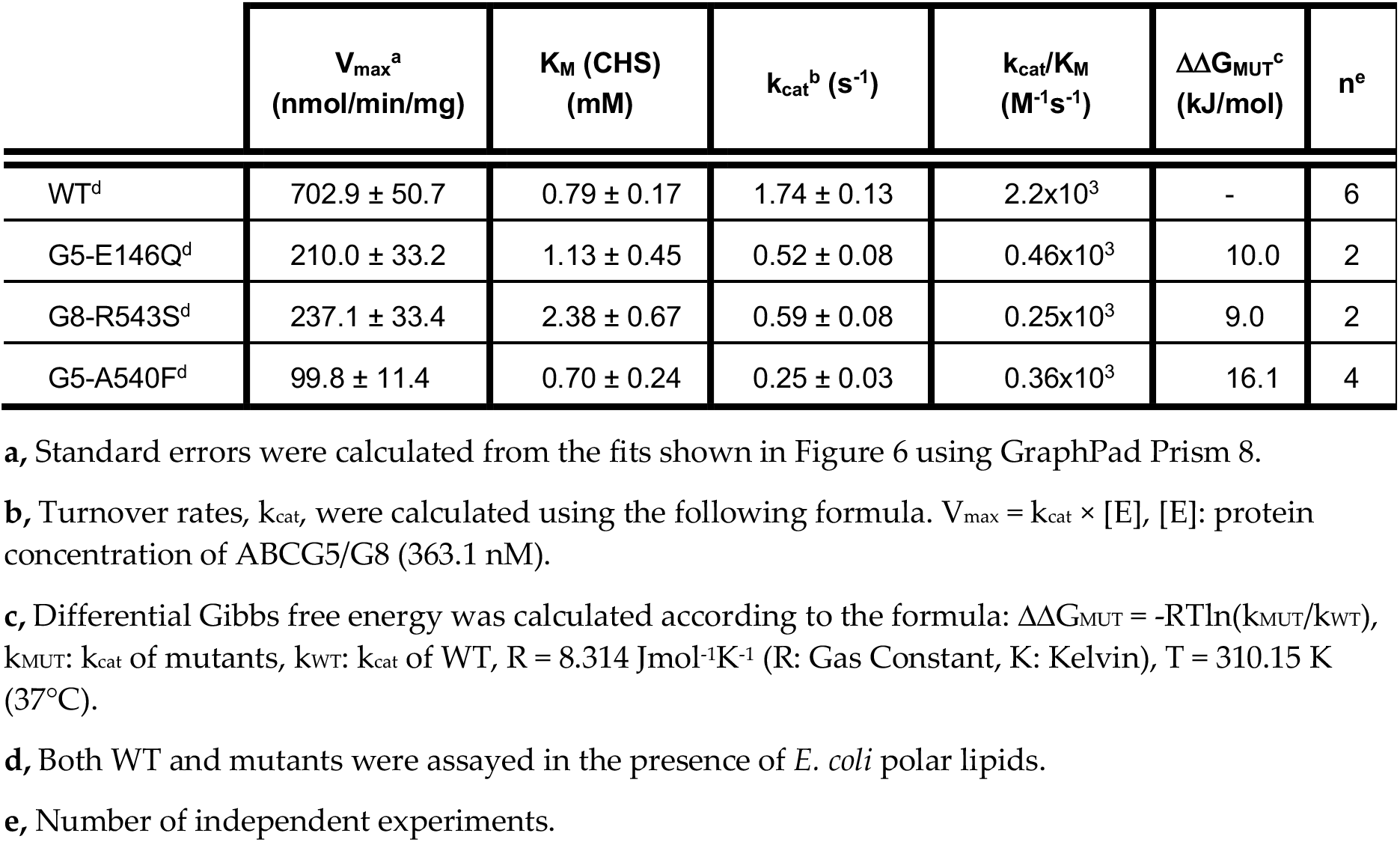
Dependence of ABCG5/G8 ATPase Activity on Cholesteryl Hemisuccinate.

| | $V_{max}^a$<br>(nmol/min/mg) | $K_M$ (CHS)<br>(mM) | $k_{cat}^b$ (s<sup>-1</sup>) | $k_{cat}/K_M$<br>(M<sup>-1</sup>s<sup>-1</sup>) | $\Delta\Delta G_{MUT}^c$<br>(kJ/mol) | $n^e$ |
| --- | --- | --- | --- | --- | --- | --- |
| WT<sup>d</sup> | 702.9 ± 50.7 | 0.79 ± 0.17 | 1.74 ± 0.13 | 2.2x10<sup>3</sup> | - | 6 |
| G5-E146Q<sup>d</sup> | 210.0 ± 33.2 | 1.13 ± 0.45 | 0.52 ± 0.08 | 0.46x10<sup>3</sup> | 10.0 | 2 |
| G8-R543S<sup>d</sup> | 237.1 ± 33.4 | 2.38 ± 0.67 | 0.59 ± 0.08 | 0.25x10<sup>3</sup> | 9.0 | 2 |
| G5-A540F<sup>d</sup> | 99.8 ± 11.4 | 0.70 ± 0.24 | 0.25 ± 0.03 | 0.36x10<sup>3</sup> | 16.1 | 4 |

**a**, Standard errors were calculated from the fits shown in Figure 6 using GraphPad Prism 8.
**b**, Turnover rates, $k_{cat}$ , were calculated using the following formula. $V_{max} = k_{cat} \times [E]$ , $[E]$ : protein concentration of ABCG5/G8 (363.1 nM).
**c**, Differential Gibbs free energy was calculated according to the formula: $\Delta\Delta G_{MUT} = -RT \ln(k_{MUT}/k_{WT})$ , $k_{MUT}$ : $k_{cat}$ of mutants, $k_{WT}$ : $k_{cat}$ of WT, $R = 8.314 \text{ Jmol}^{-1}\text{K}^{-1}$ ( $R$ : Gas Constant, $K$ : Kelvin), $T = 310.15 \text{ K}$ (37°C).
**d**, Both WT and mutants were assayed in the presence of *E. coli* polar lipids.
**e**, Number of independent experiments.

### 2.4. Missense mutations cause conformational changes at the ATP-binding site

To examine the relationship between structural changes of missense mutations and their impact on the ATPase activity, we have performed molecular dynamics (MD) simulations for the WT and three mutants in this study. We then analyzed the MD structures to understand how the mutations could lead to different conformation around the hypothetical surrounding residues at the nucleotide-binding sites (NBS). These residues were obtained through a structural comparison between the crystal structure of ABCG5/G8 (PDB ID: 5DO7) and a cryo-EM structure of ABCG2 (PDB ID: 6HBU) for which two ATP were bound in the homodimer [21,38].

To identify which residues are important for the ATP binding, we conducted MD simulations for the ABCG2 system. We calculated the ligand-residue MM-GBSA free energies (ΔG_lig-res_) for the 32 surrounding residues and identified eight hotspot residues which have ΔG_lig-res_ better than −7.0 kcal/mol (**Supplementary Table 1**). Although those hotspots were identified for ABCG2, it is reasonable to assume they are also hotspots for ABCG5/G8 given the apparent structural and sequence similarity (only one hotspot has different amino acid types). The root-mean-square deviation (RMSD) for the mainchain atoms is 2.60 Å, and the corresponding amino acid types of both proteins are listed in **Supplementary Table 1**. The detailed interactions between ATP and ABCG2 revealed by a representative MD structure is shown in **Supplementary Figure 3**. In this study, we have focused on the active nucleotide-binding site (known as NBS2) in ABCG5/G8 [21] and analyzed residues 88-103, 246-251 of ABCG5 and 210-220, 237-245 of ABCG8. Those residues were recognized as the surrounding residues of the NBS2 in ABCG5/G8.

As shown in **Figure 7**, the mutations at the three sites can lead to global changes on the overall ABCG5/G8 structure, with RMSD values larger than 2.0 Å. The difference between the RMSDs of the secondary structures is smaller, probably because more obvious changes need longer simulation time to manifest. We are especially interested in the mutational effect on the ATP-binding site and generated The RMSD *vs.* Simulation Time curves for those hypothetic surrounding residues (**Supplementary Figure 4**). We observed that the RMSDs with and without least-square (LS) fitting are very stable for the WT, whereas for G5-E146Q and G5-A540F, both the LS Fitting and No-Fitting RMSD are significantly larger. However, G8-R543S mutation did not lead to significantly larger RMSD. This is because the distance between the mutation site and ATP binding site is far away and much long MD simulations are required. Indeed, the RMSD has a trend of getting large along the MD simulation time for G8-R543S (**Supplementary Figure 4D**). We then conducted correlation analysis using an internal program to identify possible interaction pathways between the two sites. As shown in **Supplementary Figure 5**, the shortest path contains R543, E474, N155, V205 and L213. L213 is linked to four key residues for ATP binding. It is understandable that a perturbation at R543 needs long simulation time to reach the ATP binding site, given the shortest interaction path contains six residues including two ends. Overall, we observed a significant perturbation on the conformations of the putative surrounding residues due to the mutations at G5-E146Q and G5-A540F. We anticipated that G8-R543S mutation could lead to a significant conformational change at NBS2 in much longer MD simulations.

**Figure 7.**
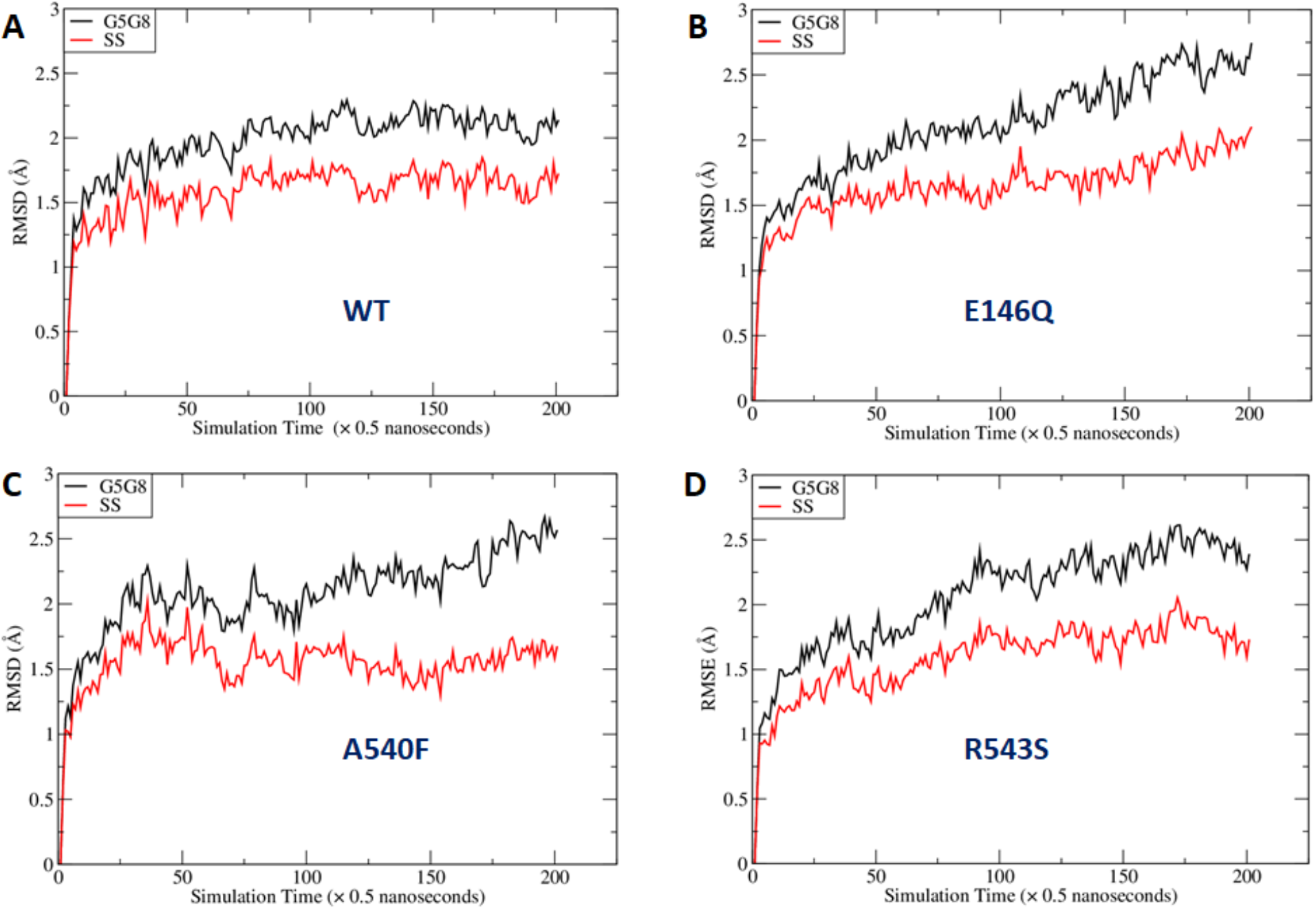
Fluctuation of Root-mean-square deviations (RMSD) along MD simulation time course. RMSD were calculated using the main chain atoms of all residues (black lines) or secondary structures only (red lines). A: wild type, B: E146Q mutant in ABCG5, C: A540F mutant in ABCG5, and D: R543S mutant in ABCG8. G5G8: ABCG5/G8; SS: secondary structure.

Next, we identified representative MD conformations for all four ABCG5/G8 protein systems for comparison (**Figure 8**). It is observed that the hotspot residues are overlaid very well between the crystal and MD structures for the WT (**Figure 8E**) and R543S mutant except for R211, while for the other two mutants, the RMSDs are significantly larger. This observation is expected, and the reason was explained above. Interestingly, the side chain of R211 underwent dramatically change for all four protein systems during MD simulations. If R211 is omitted, the mainchain RMSDs become much smaller. In summary, the conformational changes from our molecular modeling can qualitatively explain why the three mutations can lead to impaired ATPase activity. Of particular note, G5-K92, the hotspot residue that has the strongest interaction with ATP, is a part of the Walker A motif at the active nucleotide-binding site and required for ABCG5/G8 functions [18,33].

**Figure 8.**
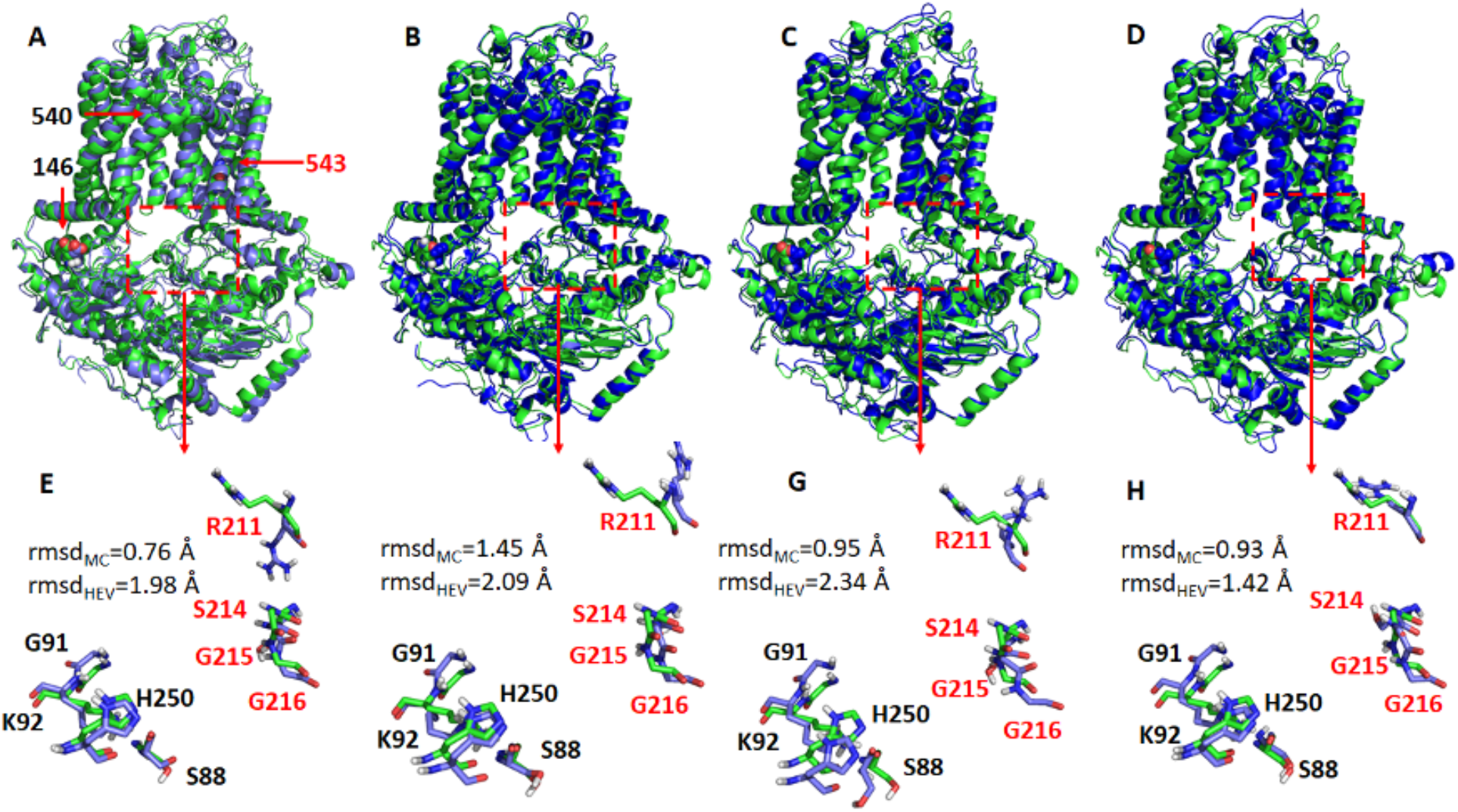
Representative structures of the WT ABCG5/G8 and its three missense mutants. The representative structures (shown as blue cartoons and bluish sticks) were aligned to the crystal structure (green cartoons, and greenish lines). The three mutation residues, E146Q, A540F and R543S, are shown as spheres. The hypothetical surrounding residues of ATP are shown as dashed rectangles. A and E: wild type, B and F: E146Q, C and F: A540F, D and G: R543S. G5: ABCG5; G8: ABCG8. Residues in G5 and G8 are separately colored in black and red. Root-mean-square deviations (RMSDs) for the mainchain atoms (rmsdMc) and all heavy atoms (rmsdHEv) were shown in the lower panels. If R211 is omitted from RMSD calculations, rmsds of the mainchain atoms are 0.69, 1.30, 0.88 and 0.78 Å for WT, E146Q, A540 and R543S, respectively; the corresponding rmsds of heavy atoms are 0.85, 1.42, 1.13 and 0.96 Å.

## 3. Discussion

In this study, we have shown that CHS stimulates the ATPase activity of the human ABCG5/G8 sterol transporter to a much higher specific activity, as compared to previously reported data. (**Tables 1 & 2**). The much increased CHS-coupled ATPase activity indicates that ABCG5/G8 may need such a high ATP catalytic rate to achieve the sterol-transport function across the cellular membranes. CHS is a relatively water-soluble cholesterol analog and is used to mimic cholesterol in membrane protein crystallization [21,39]. Our results showing CHS-stimulated ATPase activity suggest that the sterol molecules may have played a role in promoting an active conformation for the ATPase and/or enhancing the stability of ABCG5/G8. This idea of protein stability is supported by recent findings showing that CHS stabilizes a variety of human membrane proteins towards active conformations [40]. In the crystallographic study, >2% cholesterol was necessary to produce crystals capable of diffracting X-ray to better than 4 Å, and several sterol-like electron densities were suspected on the crystal structure of ABCG5/G8 [21]. Building upon previous work using bile acids [33] and androstane [32], our enzymatic results should come with no surprise that the WT ABCG5/G8 functionality and its active conformation are directly coupled with cholesterol analogs.

For ABCG5/G8-mediated ATP catalysis, we observed similar catalytic rates from the CHS and ATP concentration-dependent experiments, a V_max_ of ~700 nmol/min/mg, whereas the K_M_ values are very similar to each other, K_M_(ATP) = 0.60 mM and K_M_(CHS) = 0.79 mM (**Tables 1 & 2**). K_M_(ATP) and K_M_(CHS) can be used to implicate ATP and sterol association to the transporters during the ATP catalytic process, respectively. We therefore speculate that one ATP usage is required for sterol-protein association for one CHS (or cholesterol) molecule. Because ABCG5/G8 is believed to contain only one active NBS [18], such 1:1 stoichiometry of ATP and cholesterol for ABCG5/G8 may reflect the sterol transport rate by the single active site on this ABC transporter. An *in vitro* sterol-binding or transport assay, in need to develop, will be necessary to directly address such relationship. In addition to sterols, it is intriguing that PE, PC, or PG alone was sufficient to support ATPase activity of ABCG5/G8, with PE-driven activity the highest (**Figure 4**). PE is the major phospholipid of the *E. coli* polar lipids, ~60%, and the second abundant phospholipid in the bile canalicular membranes and the small intestine brush-border membranes, ~25% and ~40% respectively of total phospholipids [41,42]. It has been shown that PE preferentially fits the headgroup-binding sites on integral membrane proteins [43]; thus, PE may be recruited as better phospholipids to support ABCG5/G8 function in the cell membranes. The approximate ratio of lipids for either *E. coli* or liver polar lipids may contribute to the apparent difference in activity, but it remains unknown how phospholipid composition regulates the transporter function. It is worth noting that specific phospholipids were shown to regulate the ATPase activity of other ABC sterol transporters, such as sphingomyelin, although the mechanistic detail is not clear [19]. These individual lipids will be subjected to further examination to define the phospholipid specificity on the ABCG5/G8 ATPase activity and/or sterol transport function.

By mapping disease-carrying residues on the apo structure of ABCG5/G8, we have found that most missense variants occur within or near the structural motifs consisting of several conserved amino acids [22]. Several missense mutations (ER-trapped) prevent protein maturation from the endoplasmic reticulum (ER), but at least five mutations (ER-escaped) have been shown to undergo proper trafficking to post-ER cell membranes [29]. So far, no report has shown the impact of these ER-escaped missense mutants on ABCG5/G8 function using either *in vitro* or *in vivo* models. In this study, we have used purified proteins from *Pichia pastoris* to investigate the functional activity of ABCG5/G8 *in vitro* and aimed to establish the mechanistic basis of ABCG5/G8 through analyzing the structure-function relationship of its loss-of-function missense mutations. The sitosterolemia missense mutants G5-E146Q and G8-R543S have shown a reduction of CHS-coupled ATP hydrolysis, but retained ~20% activity as compared to WT, while the putative sterol-binding mutant G5-A540F has shown further reduction to ~10% of WT ATPase activity (**Figures 5 & 6**). With such activity reduction, the mutant proteins maintained the ATPase activity similar to the basal level, as shown by WT, suggesting a remote and allosteric regulation to keep ATPase active during the reaction.

It is not uncommon that reagents such as CHS may be used as protein stabilizers for disease-causing missense variants. Here, in the absence of CHS-coupled stimulation, the mutants showed similar level of reduced ATPase activity, arguing a more profound effect from impaired allosteric regulation on the catalytic activity of the mutants, rather than CHS-driven stability for mutant proteins. As predicted by MD simulation, the ATP-bound homology model underwent global conformational changes upon introducing the mutations (**Figure 7**). These mutations, albeit relatively far away from the nucleotide-binding site, can cause significant structural rearrangement of the residues within the region that encompass the active NBS2 (**Figure 8**). Such conformational changes may alter responses to sterol-protein interaction necessary for maximal ATPase activity.

In the atomic model of ABCG5/G8 (PDB ID: 5D07), G5-E146 is located on the hot-spot helix of the triple-helical bundle and in proximity to ABCG5’s polar relay, while G8-R543 is part of ABCG8’s polar relay in the core of TMD (**Figure 1**). Both triple-helical bundle and polar relay are believed to form a network of hydrogen bonding and salt bridges and play an important role in inter-domain communication during the transporter function [21]. G5-E146 and G8-R543 are found in the proximity of the hydrogen-bond distance with Arginine 377 of ABCG5 (G5-R377) and Glutamate 503 of ABCG8 (G8-503), respectively (**Figure 1B**). Based on the ATP-dependent experiments (**Figure 5 & Table 1**), we obtained the changes of Gibbs free energy from WT to each mutant (ΔΔG_MUT_) as ΔΔG_EI46Q_= ~11.7 kJ/mol and ΔΔG_R543S_= ~12.3 kJ/mol. Such energetic loss is in the range of intramolecular hydrogen-bonding potential observed on transmembrane a-helical bundles [44]. Therefore, the results support the hypothesis that the hot-spot helix and the polar relay are responsible for transmitting signals between NBD and TMD. Slightly lower ΔΔG_MUT_ was observed from CHS-dependent experiments (**Figure 6 & Table 2**), ΔΔG_EI46Q_= ~10.0 kcal/mol and ΔΔG_R543S_= ~9.0 kcal/mol. This falls in the range of hydrophobic interaction and argues weakened sterol-transporter interaction due to these disease mutations. As for the sterol-binding mutant, we obtained higher energetic loss, but similar ΔΔG_MUT_ from ATP- or CHS-dependent analysis, ΔΔG_A54OF_= ~15.8 or ~16.1 kJ/mol, respectively. This likely indicates a strong hydrophobic interaction between sterols and the transporter, as no obvious hydrogen donors/acceptors can be found at the putative sterol-binding site on the crystal structure. In addition, G5-A540 is distant from the polar relay (>10Å away); thus, this data suggests a remote contact by sterol molecules to control the sterol-coupled signaling, likely through the polar relay in the transmembrane domains. In the ATP concentration-dependent experiments, the K_M_ values for ATP remained almost the same (**Table 1**), suggesting that ATP binding was not affected by these mutants. The K_M_ values for CHS was significantly increased in the disease mutants, but not the sterol-binding mutant (**Table 2**), suggesting that CHS interacts with ABCG5/G8 and remotely regulates the turnover of ATP hydrolysis in either sequential (Mode 1) or concerted (Mode 2) pathway (**Figure 9**). Collectively, these results argue that a working network of hot-spot helix and polar relay is essential to maintain the communication between ATPase and sterol-binding activities in ABCG5/G8, which are impaired by the loss-of-function missense mutations. As G8-R532S is the only know ER-escaped disease mutant, we will expect more insight in such polar relay-driven allosteric regulation by investigating other polar relay residues with site-directed mutagenesis.

**Figure 9.**
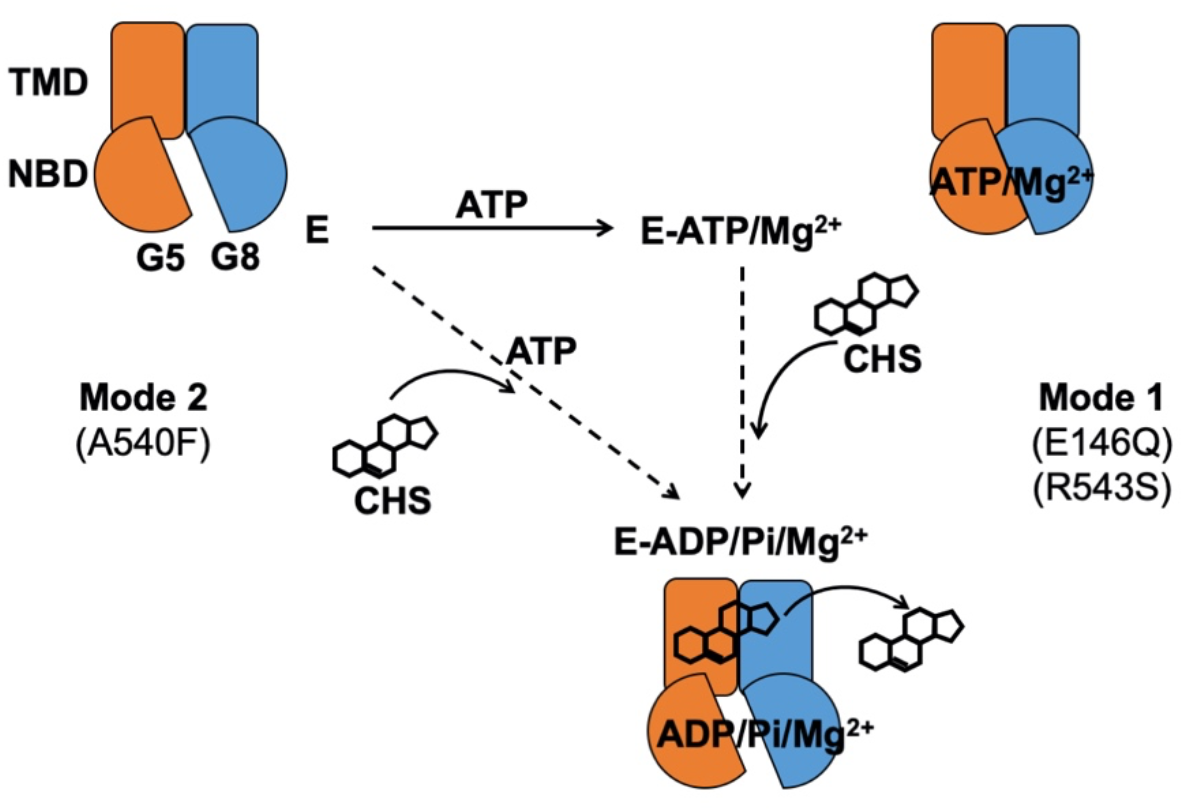
Proposed mechanism of sterol-coupled ATP catalysis by ABCG5/G8. *(Mode 1)* A sequential pathway is derived from experiments on the disease mutants, G5-E146Q and G8-R543S. ABCG5/G8 first recruits ATP and Mg^2+^ ions, likely causing a conformational change of the NBD for ATP binding. CHS/sterol then binds the transporter and triggers ATP hydrolysis that may result in its dissociation. *(Mode 2)* A concerted pathway is derived from experiments on the putative sterol-binding mutant, G5-A540F. ABCG5/G8 simultaneously recruits CHS, ATP, and Mg^2+^ ions, induces a transient conformational change of the NBD, and activates ATP hydrolysis and CHS/sterol dissociation from the transporter. G5: ABCG5; G8: ABCG8; E: ABCG5/G8 heterodimer; Pi: inorganic phosphate.

In conclusion, these studies show that CHS stimulates ABCG5/G8 ATPase activity and may promote an active conformation for ABCG5/G8-mediated sterol transport. The enzymatic characterization of three loss-of-function missense variants provides a mechanistic basis of how the polar relay contributes to the inter-domain communication for the sterol-coupled ATPase activity in ABCG5/G8 and may be directly involved in such ligand-protein interactions. Further studies will reveal more insight into these molecular events and enable sterol-lowering therapeutics to treat sitosterolemia and hypercholesterolemia.

## 4. Materials and Methods

### 4.1. Materials

*E. coli* polar lipids (Cat. #: 100600C) and bovine liver polar lipids (Cat. #: 181108C) were from Avanti Polar Lipids, Inc. (via MilliporeSigma). Cholesterol, cholesteryl hemisuccinate (CHS), and n-Dodecyl β-D-maltopyranoside (DDM) were from Anatrace. The Ni-NTA agarose resin was from Qiagen, the calmodulin (CBP) affinity resin, zeocin, and ampicillin were from Agilent. Imidazole, ε-aminocaproic acid, sucrose, yeast extract, tryptone, peptone, yeast nitrogen base (YNB), and ammonium sulfate were obtained from Wisent. ATP disodium trihydrate, Tris-(2-carboxyethyl)-phosphine (TCEP), sodium chloride, glycerol, ethylene diamine-tetraacetic acid (EDTA), ethylene glycol-bis(β-aminoethyl ether)-N,N,N′,N′-tetraacetic acid (EGTA), sodium dodecyl sulfate (SDS), Ponceau S solution, sodium azide, Bradford reagents, Tween 20, magnesium chloride, calcium chloride, and all protease inhibitors were obtained from Bioshop Canada. Biotin, sodium cholate hydrate, L-ascorbic acid, ammonium molybdate, bismuth citrate, sodium citrate, methanol, ammonium hydroxide, hydrochloric acid, and acetic acid were obtained from MilliporeSigma. Dithiothreitol (DTT), Tris base and Tris acetate were obtained from ThermoFisher. Clarity Western ECL substrates, 30 % acrylamide, agarose and ammonium persulfates were obtained from Bio-Rad. Restriction enzymes were obtained from New England Biolabs, Promega and ThermoFisher. YPD: Yeast extract Peptone Dextrose; YPDS: Yeast extract Peptone Dextrose Sorbitol; MGY: Minimal Glycerol Yeast nitrogen base; mPIB (minimal protease inhibitor buffer for yeast cell lysis): 0.33 M sucrose, 0.3 M Tris-HCl (pH 7.5), 1 mM EDTA, 1mM EGTA, and 100mM ε-aminocaproic acid, ddH2O to a final volume of 1 L and stored at 4°C.

### 4.2. Cloning of ABCG5/G8 missense mutants

The expression vectors (pLIC and pSGP18), carrying human ABCG5 (NCBI accession number NM_022436) and human ABCG8 (NCBI accession number NM_022437), were derived from pPICZB (Invitrogen) as described [33,45], pLIC-ABCG5 and pSGP18-ABCG8, respectively. A tandem array of six histidines separated by glycine (Hi_S6_GlyHi_S6_) was added to the C terminus of ABCG5, and A tag encoding a rhinovirus 3C protease site followed by a calmodulin binding peptide (CBP) was added to the C terminus of ABCG8. To generate the missense mutants in this study, we performed site-directed mutagenesis by using WT ABCG5 or ABCG8 as the templates and the following codon-optimized oligonucleotide primers (Eurofins Genomics Canada). G5-A540F: CCATTTTTGGGGTGCTTGTTGGATCTGGATTCCTCAG (forward) and GCACCCCAAAAATGGACAGCAGAGCCACTACAC (reverse); G5-E146Q: GCGCCAAACGCTGCACTACACCGCGCTGC (forward) and CAGCGTTTGGCGCACGGTGAGGCTGCTCAG (reverse); G8-R543S: GTTGCTCTATTATGGCCCTGGCCGCCGC (forward) and GCCATAATAGAGCAACAGAAGACCACCAGCCAC (reverse); G8-G216D: ACGAGCGCAGGAGAGTCAGCATTGGGGTGCAG (forward) and CTCTCCTGCGCTCGTCCCCCGACAACCCC (reverse). The polymerase chain reaction (PCR) included 1-unit Phusion High-Fidelity DNA Polymerase (New England Biolabs), 1x Phusion buffer, 200 mM dNTP, 2% (v/v) DMSO, 100 ng DNA templates, and 0.4 mM forward and reverse primers. Each mutant-containing plasmid was amplified by the following PCR setting: initial DNA denaturation (98°C, 2 minutes), followed by 30 cycles of denaturation (98°C, 15 seconds) / primer annealing (55°C, 30 seconds) / DNA extension (72°C, 3 minutes), then final extension (72°C, 20 minutes). 5μl of the PCR products was run on a 1% agarose gel to confirm the amplification, and 1μl of Dnp1 restriction enzymes (20 units) was used to digest the WT templates overnight at 37°C to. The modified plasmids were cleaned up by ethanol acetate precipitation technique. 5μl of 3M Sodium acetate was added to each 50μl PCR product. 200μl of 100% Ethanol was added to each tube, vortexed, and left at room temperature for 10 minutes. At max speed in a table centrifuge for 10 minute the plasmids were pelleted the supernatant was removed then washed by 75% ethanol. Residual ethanol was dried by a Speed-Vac at the maximal speed for 20 minutes at room temperature. The pellet was resuspended in ddH2O. Mutants plasmids were cloned into XL1-Blue competent *E. coli* cells by the heat-shock approach as described in the supplier’s manual (Novagen/Agilent) and by antibiotic selection using Zeocin (Invitrogen/ThermoFisher). Using PureYield Plasmid Midiprep kit (Promega), DNA preparations of selected clones were subjected to sequencing at Eurofins Genomics Canada.

### 4.3. Expression of ABCG5/G8 missense mutants in *Pichia pastoris* yeast (Supplementary Figure 1A)

Both WT and mutant plasmids (20 mg each plasmid) were linearized using PmeI and cotransformed into the *Pichia* strain KM71H by electroporation. Immediately the cells were resuspended with 1-mL ice cold 1M sorbitol and incubated at 30°C for 1 hour. Then 5mL fresh YPD were added and incubated for 6 hours at 250 rpm and 30°C. The cells were then centrifuged at 3000xg for 10 minutes and resuspended with 200μL of YPD. 100 μL of transformants were plated on YPDS plates containing 100 (low), 500 (medium) or 1000 (high) μg/mL of Zeocin to screen for successful transformation. Seven colonies were picked and grown in 10 mL of MGY media for 24 hours in sterile 50 mL tubes at 250 rpm and 30°C. The cells were centrifuged for 10 minutes at 3000xg and the resuspended with 10 mL of MM media. 50 μL methanol was added to the media and once again after 12 hours. The cells were harvested after 24-hour incubation at 250 rpm and 30°C, resuspended in 600 μL mPIB buffer transferred into 1.5 mL Eppendorf tube. After adding 500 μL glass bead, protease inhibitors, and 10mM DTT, the cells were lysed using a mini-bead beater (Biospec), 1.5 minutes beating and 1.5 minutes rest on ice for 3 cycles. The unbroken cells and beads were pelleted by centrifugation at 5000×g for 5 minutes at 4°C, followed by 21130xg for 5 minutes at 4°C. The supernatant was collected, and the concentration of the total proteins was estimated by Bradford assay. 1 μl of cell lysate alongside a 0 μg to 10 μg BSA standards were each added to a 200 μl Bradford reagent on a 96-well plate. Absorbance at 595 nm was used to measure the protein concentrations using a Synergy H1 Hybrid reader. The cell lysates (20 or 30 μg of total proteins) was resolved by SDS–PAGE, and protein expression was analyzed by immunoblotting using monoclonal anti-RGSH4 antibodies (Qiagen) to detect ABCG5 and polyclonal anti-hABCG8 antibodies (Novus Biologicals) to detect ABCG8. The clones expressing the highest level for both subunits were selected and stored in 20% glycerol at −75°C.

### 4.4. Cell culture and microsomal membrane preparation

The conditions for cell growth and WT protein induction were as described (14). Briefly, cells were initially grown at 30°C to accumulate cell mass in an Innova R43 shaker (Eppendorf) at 250 rpm for 24-48 hours with the pH maintained at pH 5-6. To induce protein expression, cells were left fasting for 6-12 hours, then incubated with 0.1% (v/v) methanol for 6–12 hours at 20 or 28°C. The methanol concentration was increased to 0.5% (v/v) by adding methanol every 12 hours for 48-60 hours. Cell pellets were collected and resuspended in mPIB and stored at −75°C. Approximately 45 ± 10 g of cell mass was typically obtained from 1 L of cultured cells. The frozen cells were thawed and lysed using a C3-Emulsifier (Avestin) in mPIB in the presence of 10 mM DTT and protease inhibitors (1 μg/ml leupeptin, 1 μg/ml pepstatin A, 1 μg/ml aprotinin, and 2 mM PMSF). The microsomal membranes were then prepared, as described [21].

### 4.5. Purification of ABCG5/G8 and its mutants

Both WT and mutants were purified following a protocol as described previously [21], with minor modification. Briefly, DDM-solubilized membranes were subjected to a tandem affinity column chromatography, first using Ni-NTA and then CBP. The N-linked glycans and the CBP tag remained on the purified heterodimers, and the CBP eluates were further purified by gel-filtration chromatography using a Superdex 200 Increase 10/300 GL column on an ÄKTA Pure purification system (Cytiva, formerly GE Healthcare Life Sciences). The proteins in the peak fractions were collected and concentrated to 1-3 mg/mL for storage at −75°C. Noticeably, the final yield for mutants was lower than WT, in a range of 400-800μg per six liters of cells. The expression level of the mutant proteins in the microsomes and their solubility was slightly lower than WT. Some proteins were also lost during Ni-NTA binding and imidazole wash. The profile of the gelfiltration chromatography often showed a higher peak at the void volume than dimeric proteins. These factors collectively suggest that the mutant proteins are more prone to aggregation, and thus explain the lower yields.

### 4.6. ATPase assay

We have consistently observed a strong cloudiness in the assay solution when using previous protocols, consequently resulting in low sensitivity of detecting the ABCG5/G8 ATPase activity. Because a high concentration of bile acids is required, we have reasoned that the high content of detergents, both in the assay solution and in the protein preparations, may have caused either high background upon quenching the reaction in the Malachite Green-based assay [33] or poor organic-aqueous phase separation [21]. The measurement of ATPase activity has thus become inconsistent from one protein preparation to another. To overcome this issue, we first optimized the ATPase assay by adopting a colorimetric and bismuth citrate-based approach [35], which also allows high-throughput detection of the liberated inorganic phosphate by a microplate reader. The ATPase assay was performed in a 65μl final reaction volume containing 2mg/ml *E. coli* or liver polar lipids or designated phospholipids, 1.5% sodium cholate, 0.2% (4.11mM) CHS, and 2mM DTT in Buffer A (50mM Tris/Cl pH 7.5, 100mM NaCl, 10% glycerol, 0.1% DDM). The Lipid/CHS/DTT mixture was thoroughly sonicated and preincubated with ABCG5/G8 proteins (0.3 to 1.5 μg) for 5 minutes at room temperature. The catalytically deficient G8-G216D was used as the negative control.

The enzymatic activity of ABCG5/G8 was initiated upon the addition of the 10X ATP cocktail (6.5μl) and incubated at 37°C. Aliquots (8.5μl) were removed every 2 minutes and added to the prechilled quencher wells to stop the reaction. The quencher solution was made of 5% SDS in 5mM HCl, which together with smaller reaction volume, contributed to significant reduction of cloudiness for inorganic phosphate detection. Lipid mixtures were prepared at 30mg/ml (~20mM) in Buffer A containing 7% sodium cholate. CHS stock solution (1%, w/v) was prepared in a Buffer A and 4.5% sodium cholate. 10X Mg/ATP cocktail contains 50mM ATP, 75mM MgCl2, 100mM NaN3 in a buffer containing 50mM Tris/Cl pH 7.5. To detect the liberated inorganic phosphate, 50μL of freshly-made Solution II (142mM ascorbic acid, 0.42M HCl, 4.2% Solution I (10% ammonium molybdate) was added to plate wells and left on ice for 10 minutes. Then 75μl Solution III (88mM bismuth citrate, 120mM sodium citrate, 1M HCl) was added to plate wells and placed at 37°C for 10 minutes. The absorbance was measured at 695nm using a multi-well plate reader. For the phosphate standards, 1M monobasic or dibasic sodium or potassium phosphate in 50mM Tris/Cl pH 7.5 was prepared, and six standard inorganic phosphate solutions (0μM, 12.5μM, 25μM, 50μM, 100μM, 200μM) were used in every experiment. The linear range of each reaction was used to calculate the initial rate of ATP hydrolysis. GraphPad Prism 8 was used to perform nonlinear regression and ordinary one-way ANOVA, with a P value of ≤0.05 considered significant from at least three independent experiments. The kinetic parameters were calculated by nonlinear Michaelis-Menten curve fitting using GraphPad Prism 8.

### 4.7. Computational Methods

We studied four ABCG5/G8 protein systems including the WT, E146Q, A540F and R543S mutants. Each MD system consists of one copy of ABCG5/G8 heterodimer, 320 1,2-Dimyristoyl-sn-glycero-3-phosphocholine (DMPC) lipid, 16 Cholesterol, 43,621 TIP3P [46] water molecules, 103 Cl^-^ and 83 Na^+^ to neutralize the MD systems. AMBER ff14SB [47], Lipid14 [48] and GAFF [49] force fields were used to model proteins, DMPC lipids and Cholesterols, respectively. The residue topology of cholesterol was prepared using the Antechamber module [48]. MD simulation was performed to produce isothermal-isobaric ensembles using the pmemd.cuda program in AMBER 18 [50]. The Particle Mesh Ewald (PME) method [51] was used to accurately calculate the electrostatic energies with the long-ranged correction taken into account. All bonds were constrained using the SHAKE algorithm [52] in both the minimization and MD simulation stages following a computational protocol described in our previous publication [21]. Briefly, there were three stages in a series of constant pressure and temperature MD simulations, including the relaxation phase, the equilibrium phase, and the sampling phase. In the relaxation phase, the simulation system was heated up progressively from 50K to 250K at steps of 50K and 1-nanosecond MD simulation was run at each temperature. In the next equilibrium phase, the system was equilibrated at 298K, 1 bar for 10 ns. Finally, a 100-nanosecond MD simulation was performed at 298K, 1 bar to produce isothermal– isobaric ensemble ensembles. In total, 1,000 snapshots were recorded from the last phase simulation for post-analysis using “cpptray” module implement in the AMBER software package. Binding free energy decomposition and correlation analysis were performed using an internal program and the detailed elsewhere [53,54].

## Author Contributions

B.M.X. optimized the CHS-stimulated ATPase assay for ABCG5/G8; A.A.Z. generated and validated the mutant constructs; B.M.X., A.A.Z. and A.V. purified the proteins and carried out the ATPase assays and data analysis. J.W. performed the molecular dynamics simulation, J.-Y.L. oversaw the project, and J.W. and J.-Y.L. wrote the manuscript. All authors have read and agreed to the submitted version of the manuscript.

## Acknowledgments

We thank William Jennings, Gloria Ihirwe, Midhet Hajira and Chloé van de Panne for technical assistance. We also thank Donna Clary and Hui Li for their technical help in utilizing the common core facilities. We are indebted to critical feedback on reviewing and editing the manuscript from Ms. Vicky Brandt, and Drs. John Baenziger, Jean-François Couture, Gregory Graf and Xiaohui Zha. This work is partially based on the theses that were submitted to fulfill in part the requirement for the degrees of Master of Science (A.A.Z.) and Honors Bachelor of Science (A.V.).

## Abbreviations

ABC: ATP-binding cassette
ABCC7: ATP-binding cassette sub-family C member 7
ABCG5: ATP-binding cassette sub-family G member 5
ABCG8: ATP-binding cassette sub-family G member 8
ATP: Adenosine triphosphate
CBP: Calmodulin-binding peptide
CFTR: Cystic fibrosis transmembrane conductance regulator
CHS: Cholesteryl hemisuccinate
DDM: Dodecyl maltoside or n-Dodecyl β-D-maltopyranoside
DMPC: 1,2-Dimyristoyl-sn-glycero-3-phosphocholine
DNA: Deoxyribonucleic acid
DTT: Dithiothreitol
EDTA: Ethylene diamine-tetraacetic acid
EGTA: Ethylene glycol-bis(β-aminoethyl ether)-N,N,N′,N′-tetraacetic acid
ER: Endoplasmic reticulum
ICL: Intracellular loop
LDL: Low-density lipoprotein
LOF: Loss of function
LS: Least square
MD: Molecular dynamics
MGY: Minimal glycerol yeast nitrogen base
MM: Minimal methanol
mPIB: Minimum protease inhibitor buffer
NBD: Nucleotide-binding domain
NBS: Nucleotide-binding site
Ni-NTA: Nickle-nitrilotriacetic acid
PC: Phosphatidylcholine
PCR: Polymerase chain reaction
PDB: Protein data bank
PE: Phosphatidylethanolamine
PG: Phosphatidylglycerol
RCT: Reverse cholesterol transport
RMSD: Root mean square deviation
SDS: Sodium dodecyl sulfate
TCEP: Tris-(2-carboxyethyl)-phosphine
TICE: Transintestinal cholesterol efflux
TMD: Transmembrane domain
WT: Wild type
YNB: Yeast nitrogen base
YPD: Yeast extract peptone dextrose
YPDS: Yeast extract peptone dextrose sorbitol

## Appendix A

**Supplementary Figure 1.**
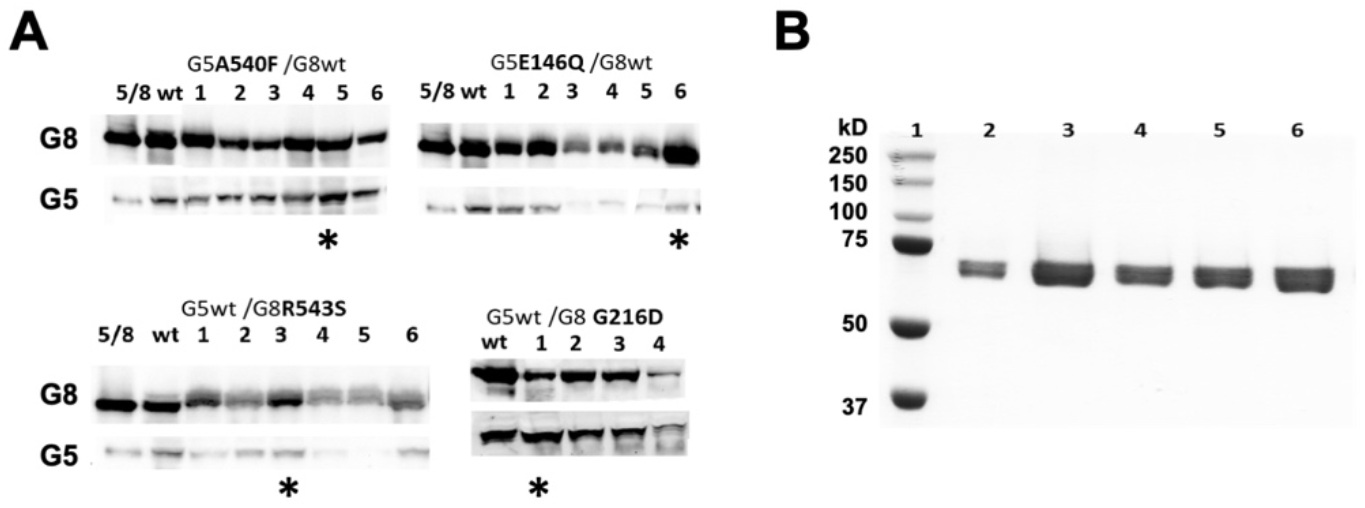
Expression and purification of ABCG5/G8 missense mutants. **A**, Four or six yeast colonies were selected for expression test. Crude microsomal membranes (either WT or mutants), containing 20-30μg total proteins, were resolved by SDS-PAGE. Protein expression was analyzed by Western blotting using a monoclonal anti-RGSH4 antibody to detect ABCG5 and a polyclonal anti-human ABCG8 antibody to detect ABCG8. The clones expressing the highest level for both subunits were selected for protein purification. Selected clones are indicated as asterisks. **B**, Gel-filtration purified mutants were resolved on a 10% SDS-PAGE gel and stained by Coomassie Blue (shown here in greyscale). Lanes 1: molecular weight marker, 2: G8-G216D, 3: G5-E146Q, 4: G8-R543S, 5: G5-A540F, and 6: WT.

**Supplementary Figure 2.**
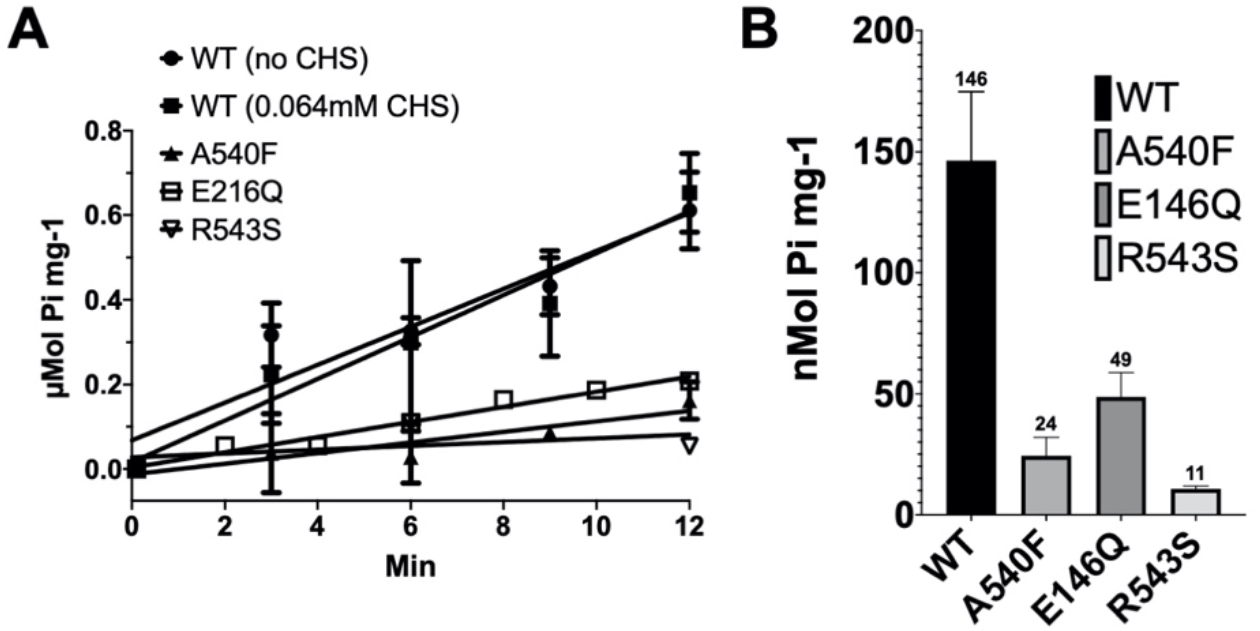
Non-CHS-stimulated ATPase activity of ABCG5/G8. **A**, The ATP hydrolysis by WT or mutant ABCG5/G8 was measured at 37°C in presence of 5mM ATP and 0.064mM CHS, a condition that resulted in consistent measurement of mutant-mediated ATPase activity and that has no appreciable effect on the WT-mediated ATPase activity. An assay protocol is described in Materials and Methods. The data points are presented as the means ± standard deviations of duplicated or triplicated experiments by using 2-4 independently purified protein preparations, where not visible, the error bars are covered by the plot symbols. A linear regression, plotted from the first 12 minutes, is used to calculate the specific activities. **B**, Bar graphs show the specific activities of non-CHS-stimulated ATP hydrolysis by WT and mutants. A540F: sterol-binding mutant G5-A540F; E146Q: sitosterolemia mutant G5-E146Q; R543S: sitosterolemia mutant G8-R543S.

**Supplementary Figure 3.**
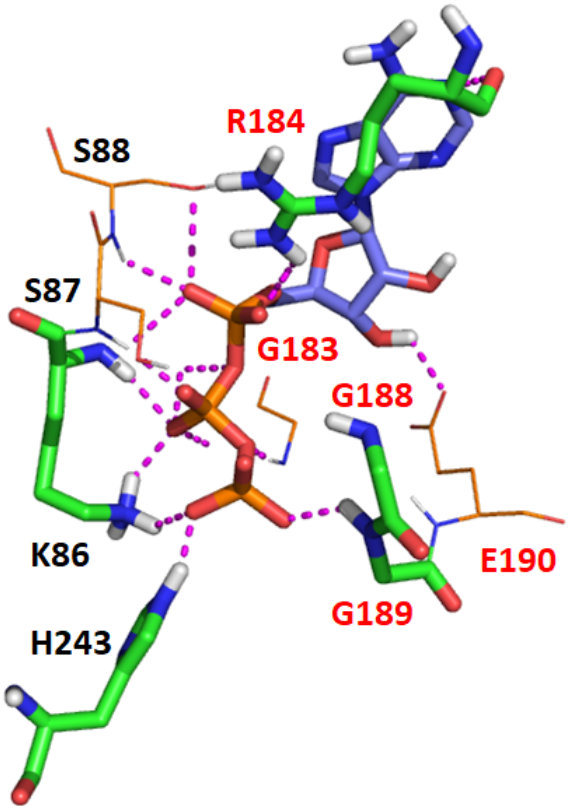
The interaction between ATP and ABCG2 revealed by a representative MD snapshot. ATP is shown as bluish sticks and hotspot residues in greenish sticks (ΔG_lig-res_ ≤ −10.0 kcal/mol). Other residues forming hydrogen bonds (dashed magenta lines) with ATP are shown in lines.

**Supplementary Figure 4.**
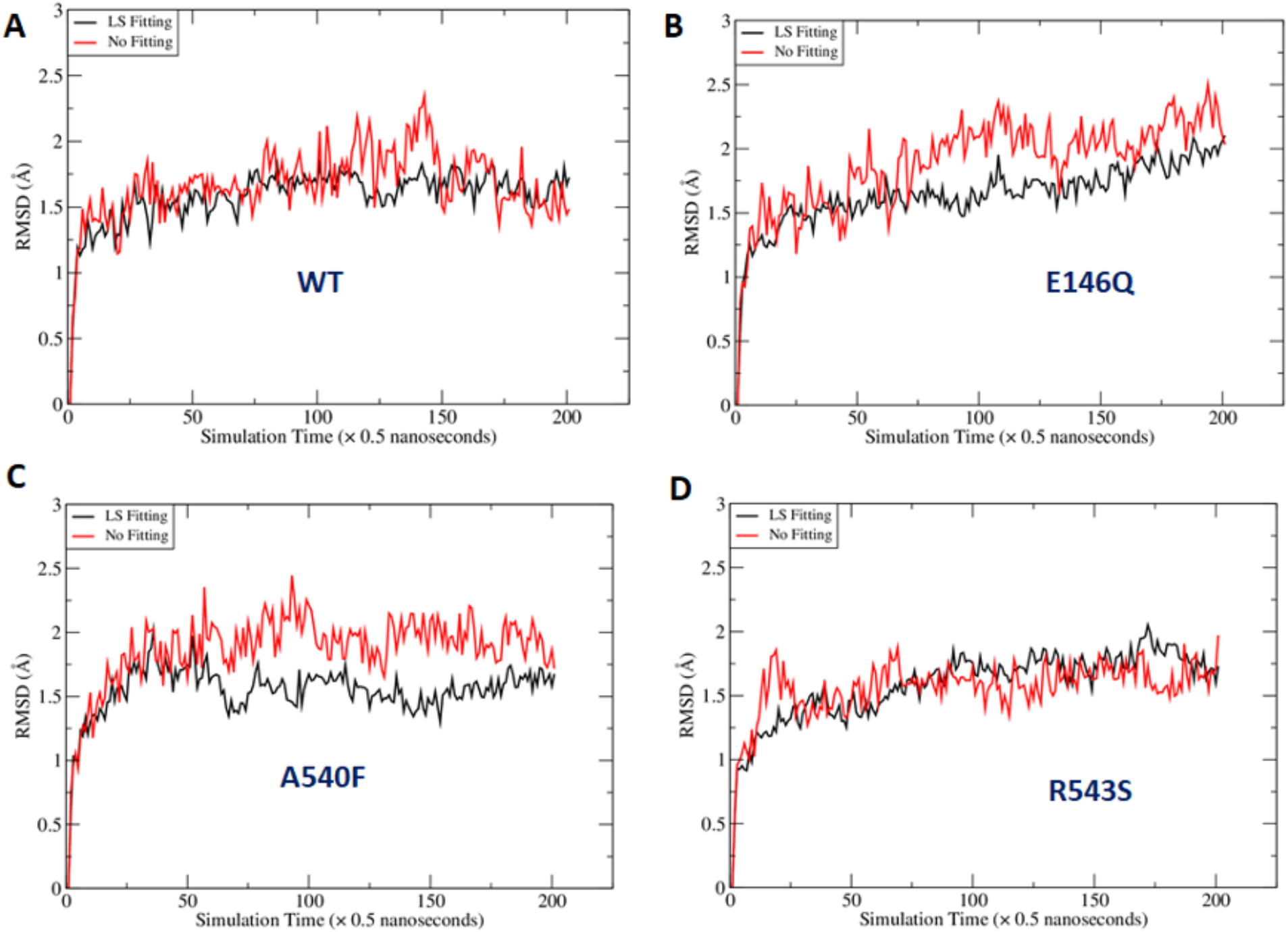
Fluctuation of Root-mean-square deviations (RMSD) along MD simulation time course. RMSD were calculated for the hypothetical residues surrounding ATP under two scenarios. In the first scenario, least-square (LS) fittings were performed for the main chain atoms of the hypothetic residues which including Residues 88-103, 246-251 of G5 and 210-220, 237-245 of G8. In the second scenario, the RMSD were calculated directly for the hypothetical residues without LS Fitting after the MD snapshots were aligned to the crystal structure using the secondary structures of ABCG5/G8. The fluctuations of the first and second scenarios were illustrated using black and red curves.

**Supplementary Figure 5.**
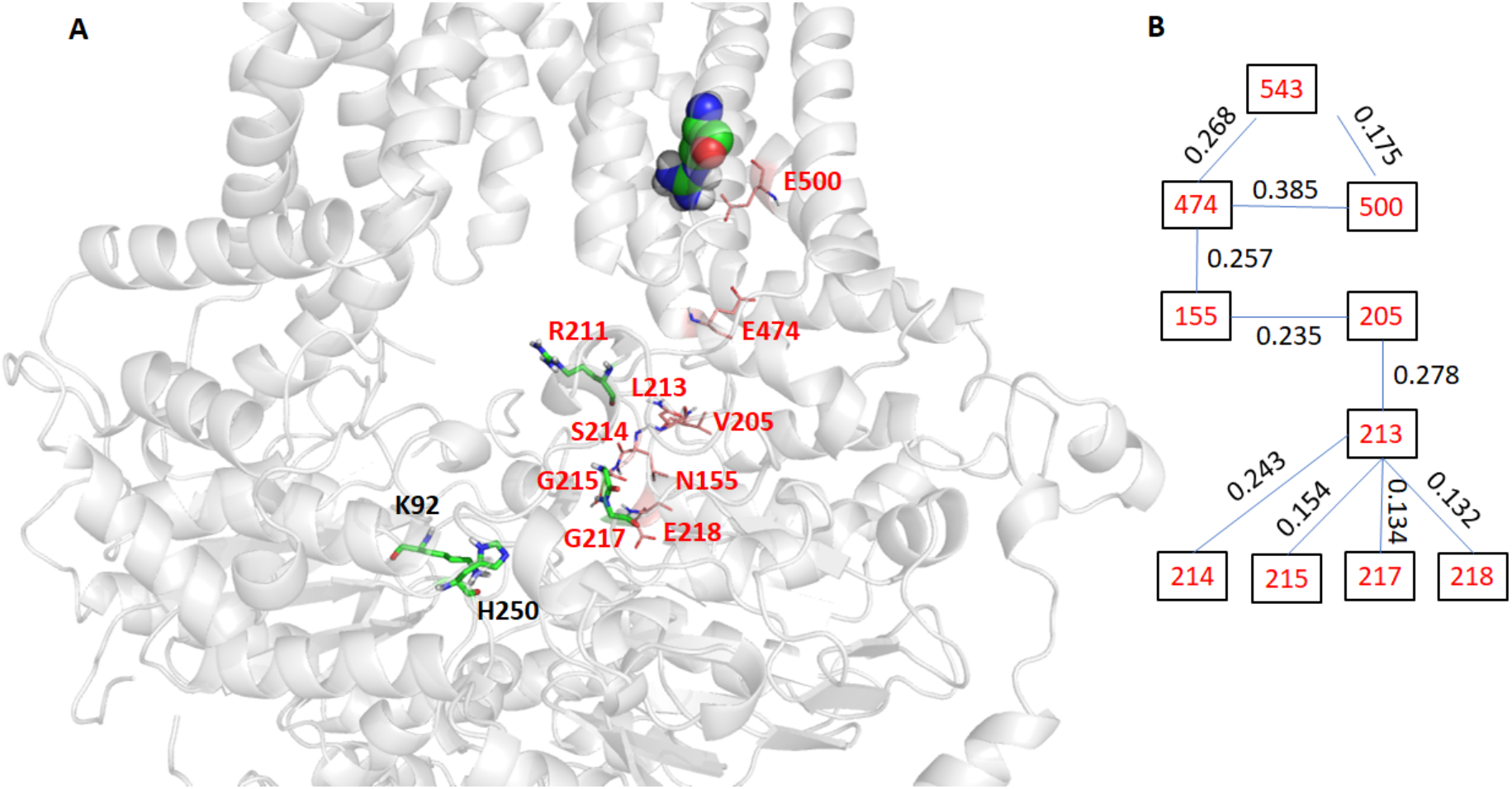
Interaction pathways link Residue R543 of ABCG8 and hotspot residues. The hotspot residues which have ligand-residue interaction energies more potent than −10.0 kcal/mol are shown in sticks, and residues are within the interaction pathways are shown in lines and labeled in red (Panel A). The residues within the interaction pathways and the correlation between them are shown in Panel B. A correlation between two residues, which is between 0 and 1, was obtained through correlation analysis. The average correlation for all wild-type ABCG5/G8 pairs is 0.0018 and the minimum and maximum are 0 and 0.52, respectively.

**Supplementary Table 1.**
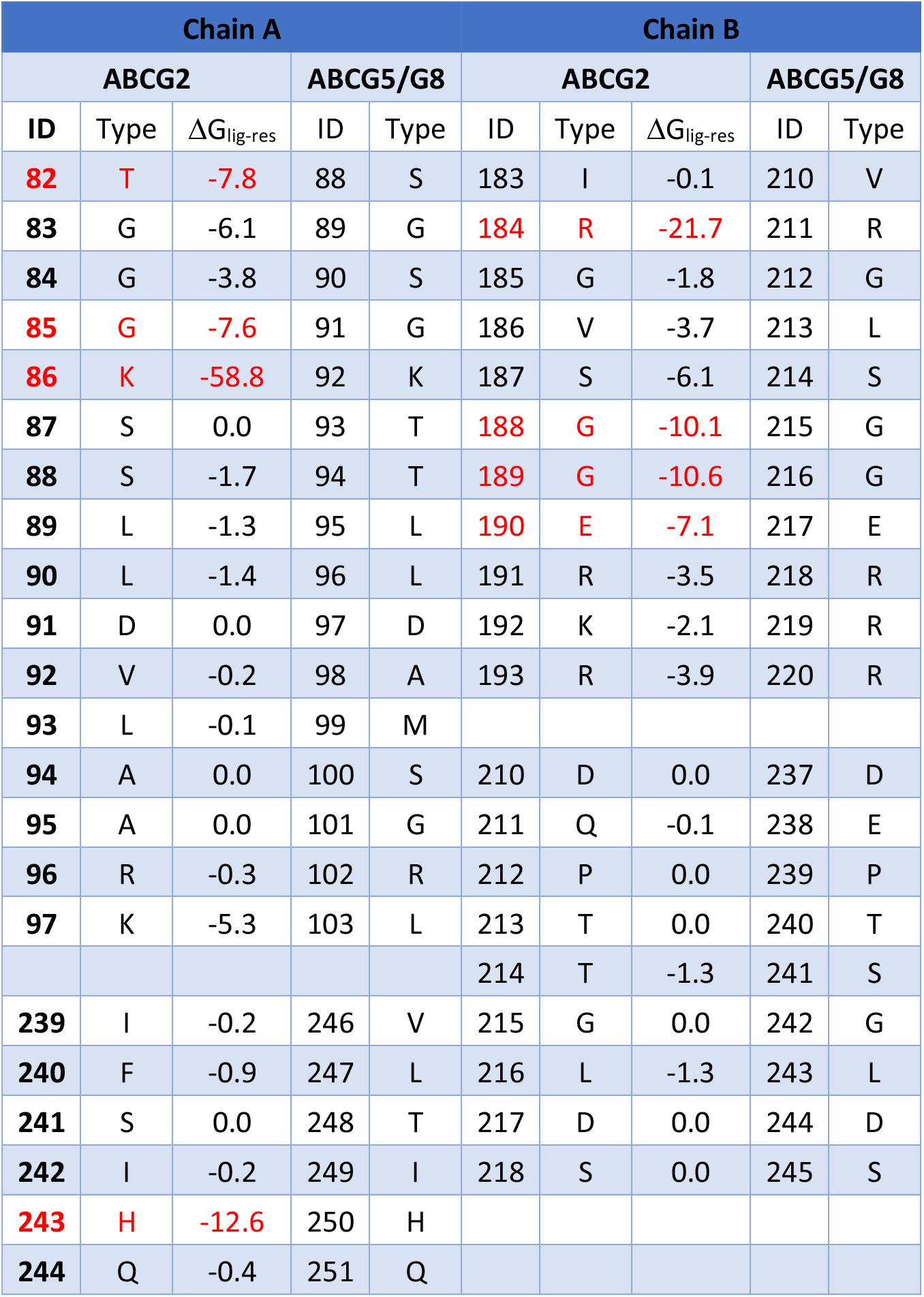
List of the hotspot residues for ABCG2 and the corresponding residues in ABCG5/G8. Hotspot residues that have ligand-residue MM-GBSA energies smaller than −7.0 kcal/mol, are shown in red.

